# Psilocybin exerts differential effects on social behaviour and inflammation in mice in contexts of activity-based anorexia (ABA)

**DOI:** 10.1101/2025.10.14.682467

**Authors:** Sheida Shadani, Erika Greaves, Zane B Andrews, Claire J Foldi

## Abstract

Psychedelics, particularly psilocybin, have shown therapeutic potential across several psychiatric conditions, including depression, anxiety, obsessive-compulsive disorder, and anorexia nervosa (AN). These disorders often share social deficits that may be effectively alleviated by psychedelics considering their use has been linked with emotional empathy and enhanced social cognition. However, the mechanisms through which psychedelics alter social behaviour are unclear, and mechanistic studies in animal models have largely focused on male subjects. This is problematic for understanding the therapeutic effects relevant for disorders that predominantly affect females, such as AN.

Here, we used the activity-based anorexia (ABA) mouse model to examine the effects of a single psilocybin dose on social behaviour in female mice and compared outcomes to mice exposed to food restriction (FR), exercise (RW) or standard housing (Controls). Together with these metabolic stressors, we also investigated the effects of psilocybin on the circulating proinflammatory cytokine interleukin-6 (IL-6), which is implicated in AN and is suppressed by psychedelics.

Psilocybin did not alter sociability in ABA, RW, or FR mice but increased preference for familiarity in Controls. Novelty-seeking behaviour was elevated in both ABA and RW groups, although with distinct social patterns. Psilocybin elevated IL-6 levels in RW mice, which was positively correlated with preference for novelty. No such relationships were found in ABA or FR groups.

These findings reveal subtle, context-dependent effects of psilocybin on social behaviour and inflammation in female mice, highlighting the need to clarify its temporal, neuroplastic, and immune-related mechanisms across sexes and disease models.

## Introduction

Anorexia nervosa (AN) is a life-threatening psychiatric condition, characterised by an inability to maintain a healthy body weight due to restrictive eating patterns, inadequate nutrition, compulsive or excessive exercise and an intense fear of weight gain (Association, 2013; Yao et al., 2016). Individuals with AN frequently exhibit perceptual distortions regarding their body weight and shape, leading to severe weight loss. Additionally, there has been a marked rise in overnight hospitalisation rates for AN among young females aged 15 to 29 in Australia, with this demographic accounting for 95% of all AN-related hospital admissions (Nguyen et al., 2022). Growing evidence suggests that AN is associated with significant impairments in emotional and social functioning, potentially stemming from deficits in social cognitive processes (Tauro et al., 2022). Social cognition is broadly defined as the set of mental processes that facilitate social interactions (Bora & Köse, 2016), enabling the interpretation of others’ thoughts, emotions and intentions (DeJong et al., 2013), which is fundamental to forming and maintaining relationships. Impaired social cognition in AN results in fewer social networks and social anhedonia (a lack of pleasure derived from social interactions) and is more pronounced in females (Harrison et al., 2014; Tauro et al., 2022). Females with AN frequently exhibit early-emerging social challenges, such as heightened feelings of loneliness, social withdrawal and perceived inferiority (Troop & Bifulco, 2002). These difficulties often precede diagnosis, with emotional empathy further diminished during the acute (weight losing) phase of AN (Morris et al., 2014). Emotional empathy refers to the capacity to recognise and share the emotional experiences of others and is a core component of social cognition (Singer, 2006). Considering the important role of empathy in prosocial behaviour, effective social functioning, interpersonal relationships and psychological wellbeing, deficits in this domain contribute substantially to the burden of AN.

Impaired social functioning is also a characteristic of other psychiatric conditions commonly comorbid with AN such as depression, anxiety disorders, and obsessive-compulsive disorder (OCD) (Godart et al., 2002; Riddle et al., 2023; Saris et al., 2017). Importantly, these conditions are consistently associated with immune dysregulation, particularly elevated pro-inflammatory signalling, reflecting a chronic neuroinflammatory state (Capuron & Miller, 2011; Lima Giacobbo et al., 2019). Pro-inflammatory cytokines can alter behaviour by modulating the metabolism of key neurotransmitters, including monoamines and glutamate, which are closely linked to stress-related symptoms such as depressed mood and anhedonia (Haroon et al., 2012). In line with this, dysregulated inflammatory responses have been implicated in the pathophysiology of depression and anxiety, with elevated levels of cytokines, particularly tumour necrosis factor-alpha (TNF-α) and interleukin-6 (IL-6), contributing to the development and persistence of these conditions (Köhler et al., 2017; Ravi et al., 2021; Renna et al., 2018). Furthermore, the potential involvement of immune dysregulation in AN is underscored by the increased frequency of infections observed in affected individuals (Gibson & Mehler, 2019). Supporting this, several studies have reported elevated levels of pro-inflammatory cytokines, particularly TNF-α and IL-6, in patients with AN (Dalton, Bartholdy, et al., 2018; Solmi et al., 2015). IL-6 plays a critical role in cognitive function and elevated IL-6 levels have also been observed in individuals with depression (Gruol, 2015; Hole et al., 2025).

One common pathway linking these features is dysfunction of the serotonin (5-HT) system, which not only regulates mood and cognition but also shapes social behaviour and modulates inflammatory responses (Bevilacqua et al., 2024; Kaye, 2008; Kiser et al., 2012; Kraus et al., 2017; Lima Giacobbo et al., 2019; Songtachalert et al., 2018). Activation of 5-HT receptors (5-HTRs) induces significant anti-inflammatory effects both centrally and peripherally, modulating immune responses in a tissue- and receptor-specific manner (Cadirci et al., 2013; Costa et al., 2020; Herr et al., 2017; Quintero-Villegas & Valdés-Ferrer, 2020). In this respect, serotonergic psychedelics that bind to 5-HTRs represent an interesting point of convergence, considering the anti-inflammatory effects of dimethyltryptamine (DMT) are well-documented (Dakic et al., 2017; Kelley et al., 2022; Nardai et al., 2020). An intriguing prospect is that psychedelics may alleviate behavioural symptoms of social dysfunction in patients via modulation of both neuronal and immune-related processes.

Among the psychiatric conditions trialled for the potential therapeutic outcomes of psychedelics, the efficacy of psilocybin (the psychoactive ingredient of magic mushrooms) in alleviating core symptoms of treatment-resistant depression is well-supported (Carhart-Harris et al., 2016; Goodwin et al., 2023). The positive therapeutic effects of psychedelics on symptoms of these conditions (including associated behavioural impairments) have been shown to include increased emotional empathy and enhanced social cognition in individuals with major depression (Bhatt & Weissman, 2024; Jungwirth et al., 2024), along with persisting prosocial outcomes, as well as anxiolytic and antidepressant effects (Carhart-Harris et al., 2018; Carhart-Harris et al., 2016; Davis et al., 2021; Goodwin et al., 2023; Griffiths et al., 2016; Grob et al., 2011; Ross et al., 2016). More recently, findings from a Phase 1 trial investigated the feasibility of psilocybin treatment in individuals with AN, indicating that the compound is safe and well-tolerated in this population (Peck et al., 2023). However, only 40% of participants exhibited significant reductions in eating disorder-related symptoms, highlighting the need to identify factors driving variability in treatment response and to clarify their neurobiological mechanisms. These gaps underscore the critical role of preclinical studies in elucidating the neurobiological pathways underlying the therapeutic effects of psychedelics.

Preclinical studies offer further evidence that psychedelics impact social behaviours in rodent models. In male mice, repeated (but not single) administration of lysergic acid diethylamide (LSD) and psilocybin enhanced social interaction and novelty preference (De Gregorio et al., 2021; Gattuso et al., 2025; Markopoulos et al., 2021) whereas a single dose of improved social reward learning (Nardou et al., 2023). However, more recent findings from a large multi-institutional study failed to replicate these effects, in which psilocybin did not enhance social reward learning or social preference in mice (Lu et al., 2025). While studies testing effects of psilocybin in animal models of psychiatric conditions are still limited, reported behavioural and neurobiological outcomes are frequently attributed to mechanisms involving synaptic plasticity (Weiss et al., 2025), but there is also evidence that inflammatory mediators of effects. Among inflammatory markers, the suppression of TNF-α and IL-6 by psilocybin and other psychedelics via 5-HT2AR activation is implicated in their antidepressant effects (House et al., 1994; Mason et al., 2023; Nau et al., 2013; Szabo et al., 2014). Given their influence on neural and immune mechanisms, the potential dual action of psychedelics on neuroplasticity and immune modulation remains an underexplored avenue of research, which could help explain their broad therapeutic potential across diagnostic categories.

To fully understand these therapeutic effects, a multifaceted approach is essential, particularly for complex psychiatric conditions like AN, which involve multifactorial mechanisms. In the present study, we used the activity-based anorexia (ABA) model to disentangle the effects of psilocybin on social behaviours and immune function in contexts of food restriction and exercise, both core features of AN. ABA is a widely recognised preclinical model that consistently captures elements of AN such as starvation-evoked hyperactivity, voluntary food restriction, severe weight loss and elevated anxiety-like behaviour (Aoki, 2021; Dong et al., 2025). Cognitive flexibility impairments in ABA rodents parallel those seen in individuals with AN (Allen et al., 2017; Huang et al., 2023) and psilocybin was recently shown to both support body weight maintenance in the ABA rat model and significantly improve cognitive flexibility (Conn et al., 2024). Building on this, we sought to understand whether psilocybin can similarly ameliorate the social and inflammatory features of AN in ABA mice, with a view to provide mechanistic insights into its potential therapeutic value for this challenging disorder.

## Materials & Methods

### Animals and housing

All animals were sourced from the Monash Animal Research Platform (MARP; Clayton, VIC, Australia). To investigate the direct effects of psilocybin on the development of the ABA phenotype, female C57Bl/6 mice (n=56), aged seven weeks upon arrival at the laboratory, were used. Young female mice were chosen for these studies due to their particular susceptibility to developing the ABA phenotype, a characteristic that remains partially understood but has translational relevance to the higher incidence of AN in young women (Nguyen et al., 2022). In all cases, animals were group-housed and acclimated to a 12-hour light/dark cycle (lights off at 0930 h) for 7 days in a temperature (22-24°C) and humidity (30-50%) controlled environment before the start of experiments. Given that the behavioural aspects of ABA (i.e., wheel running and food intake) are known to fluctuate with the estrous cycle in female mice, two male mice were placed in each experimental room at least seven days before the start of experiments to facilitate cycle synchronization, known as the Whitten Effect (Whitten et al., 1968). All experimental procedures were conducted following the Australian Code for the care and use of animals for scientific purposes and were approved by the Monash Animal Research Platform Ethics Committee (ERM 30852).

### Pharmacological compounds

Psilocybin (USONA Institute Investigational Drug Supply Program; Lot# AMS0167) was dissolved in saline and administered at a dose of 1.5 mg/kg. The compound was delivered intraperitoneally with a 26-gauge needle at an injection volume of 10 ml/kg.

### Activity-Based Anorexia (ABA)

The activity-based anorexia (ABA) model involved providing mice with the combination of unrestricted (voluntary) access to a running wheel and time-limited food access. At eight weeks old, mice were housed individually in transparent chambers equipped with a removable food basket and low-profile wireless running wheels (Med Associates Inc., USA). Mice were acclimated to the running wheel for 7 days (habituation period) to establish baseline levels of running wheel activity (RWA), which was monitored using the Wheel Manager Software (SOF-860, Med Associates Inc, USA). During the ABA period, food access was restricted to 120 minutes per day, coincident with the onset of the dark cycle (0930–1130 h). Food restriction continued until the mice reached 75-85% of their body weight from baseline. After the two-hour feeding period, the wheels were removed, and one hour later, mice were administered either saline (SAL) vehicle or psilocybin (PSI). Three control groups were included to account for the different components of the ABA model; a food restricted (FR) group that had a matched window of food access without a wheel, a running wheel (RW) group maintained on *ad libitum* food access and a singly housed control group (Control) without a wheel and with *ad libitum* access to food. Animal welfare was monitored daily, with predefined humane endpoints that included weight loss exceeding 25% of baseline.

### 3-Chamber Social Preference & Novelty Test

Mice remained in their home cage for 4-5h post-administration before being subjected to the 3- chamber social preference and novelty test (Moy et al., 2004). The test began with mice placed in the central chamber of an apparatus containing three chambers, each with a wire cage at either end, for a 10-minute habituation period. A one-minute inter-trial interval followed, during which the mice were confined to the middle chamber to minimize experimenter interference. Subsequently, an unfamiliar, age- and sex-matched mouse (novel mouse 1) was placed in a wire cage in one side chamber, while a novel object in a wire cage occupied the opposite chamber. The test mouse was then allowed to explore for 10 minutes (social preference trial). After another one-minute interval, a second unfamiliar age- and sex-matched mouse (novel mouse 2) replaced the novel object, with novel mouse 1 remaining as the familiar mouse. The test mouse was observed for another 10 minutes (social novelty trial). Locomotor activity, interaction frequency, time spent in each chamber, and social preference were measured using video tracking and EthoVision software (Noldus XT; Netherlands). Sociability was calculated using the formula: (T_s_ - T_ns_) / (T_s_ + T_ns_), where T_s_ is time spent sniffing the novel mouse, and T_ns_ is time spent sniffing the empty cage or familiar mouse. To achieve a comprehensive evaluation of behavioural metrics, emotionality-related data were normalised using Z-score methodology, as described previously (Guilloux et al., 2011). In brief, the behavioural parameters for each mouse were calculated using the following formula: *Sociability Z Score = (x - µ)/σ*

μ and σ, respectively, represent the mean and standard deviation of the singly housed (Control) group, with x referring to the individual data points for mice in the other experimental groups. The Sociability Z-Score for each mouse was determined based on both the duration and frequency of chamber entries. Sociability Z-score reflects time and frequency spent in social vs. neutral chamber, normalised to the Control group; SI reflects relative time spent sniffing the novel vs. empty cage or a familiar mouse.

### Blood biochemistry

Blood samples were collected via cardiac puncture 7-8 hours after mice had been administered PSI or SAL (control). Samples were drawn into EDTA-coated Microvette tubes (Brand) and centrifuged at maximum speed for 15 minutes at 4°C within 60 minutes of collection. Plasma was then separated and stored at -20°C until further analysis. IL-6 levels were measured using mouse IL-6 ELISA kits (KMC0061, ThermoFisher Scientifics, USA) following the manufacturer’s protocol.

ELISA data were analysed using the *Cisbio* analysis platform. Although BioSS is commonly recommended for this type of analysis, preliminary comparisons showed that BioSS flagged a significant proportion (>50%) of data points, potentially due to the stringency of its statistical thresholds or assumptions about signal uniformity. In contrast, *Cisbio* provided more consistent results across plates, with fewer flagged values and better alignment with expected standard curve behaviour. As such, *Cisbio* was selected for data processing and quantification due to its robustness and reproducibility in this context.

### Statistical analysis

All statistical analyses were conducted using GraphPad Prism version 9.5.1 (GraphPad Software, San Diego, CA, USA). A threshold for statistical significance was established at p□<□0.05, with values between p□<□0.10 considered as indicative of a trend, though not statistically significant. Data are presented as mean ± SEM and depending on the nature of the data, the number of groups, and the specific comparisons required, various analyses were employed, including two-tailed unpaired t-tests, one-way and two-way analyses of variance (ANOVA) with post hoc tests (Šidák or Tukey), as well as mixed-effects models. Outcomes of all statistical test are presented in **Supplementary Materials**.

## Results

### Establishing the ABA model in female mice

We first established the ABA model (schematic in **Figure 1A**) to investigate the effects of food restriction combined with *ad libitum* running wheel access on sociability in female mice. Throughout the habituation (wheel training) period, ABA mice had body weights comparable to the FR group. However, after 3-4 days of ABA exposure, ABA mice exhibited significantly lower body weights (Day 3: *p* = 0.036; Day 4: *p* = 0.0263; **Figure 1B**) and greater overall body weight loss compared to the FR group during the ABA period (*p* < 0.0001; **Figure 1C**). The absolute daily food intake did not differ between ABA and FR mice during habituation or ABA exposure (**Figure 1D**); however, the mean daily food intake of the ABA mice was significantly higher than the FR group during the habituation phase (*p* = 0.0114; **Figure 1E**), with no difference observed during ABA exposure.

**Figure 1.**
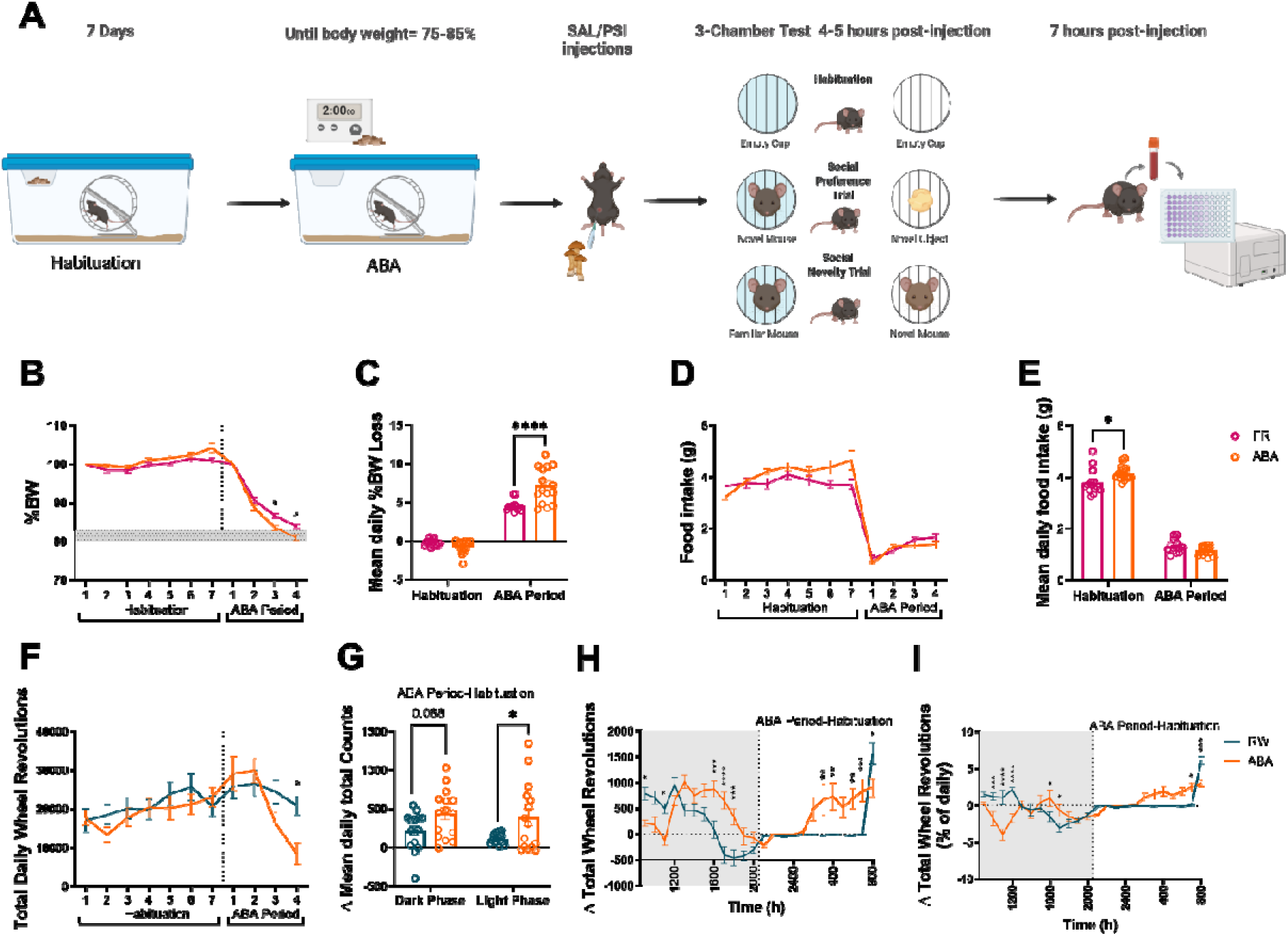
ABA elicits exacerbated weight loss and excessive exercise in mice compared to FR and RW controls. (A) Schematic of the experimental protocol. (B) Body weights were comparable between ABA and FR groups during habituation but decreased more rapidly in ABA mice demonstrated by significantly lower %BW on Day 3 (p = 0.036) and Day 4 (p = 0.0263). (C) Mean daily body weight loss of ABA mice was significantly greater compared to FR mice during the ABA period (p < 0.0001). (D) Food intake was comparable between ABA and FR groups during both habituation and the ABA phase however, (E) mean daily food intake revealed a higher food consumption in ABA mice compared to FR mice during the habituation period (p = 0.0114). (F) ABA and RW mice displayed similar running activity across periods, with the exception of Day 4, when ABA mice ran less (p = 0.0245). (G) ABA mice showed a trend towards higher running activity during the dark phase (p = 0.088) and significantly more activity during the light phase (p = 0.0294) compared to RW controls. (H) During the ABA phase, ABA mice ran less than RW mice during the food-access window (0900h: p = 0.0477; 1100h: p = 0.0176), but more post-prandially (1600h: p = 0.0006; 1700h: p < 0.0001; 1800h: p = 0.0005) and during the light phase (0300h: p = 0.0043; 0400h: p = 0.0018; 0600h: p = 0.0026; 0700h: p = 0.0001), while running less at 0800h (p = 0.01). (I) Compared to RW mice, ABA mice ran less during food access (1000h: p = 0.001; 1100–1200h: p < 0.0001) and late in the light phase (0800h: p = 0.0004), but more post-prandially (1600h: p = 0.0136; 1700h: p = 0.0301). Data are presented as mean ± SEM. Statistical analyses were performed using mixed-effects models or two-way ANOVA with Šidák post hoc tests. ABA (Activity-Based Anorexia), FR (food-restricted), RW (running wheel) groups. Significance levels are indicated as: * p < 0.05; ** p < 0.01; *** p < 0.001; **** p < 0.0001.

Similarly, while activity did not differ between ABA and RW groups during habituation; on Day 4 of the ABA phase, ABA mice exhibited significantly lower activity (*p* = 0.0245; **Figure 1F**) which was likely related to their declining body weight. Analysis of the change in daily RWA from habituation to ABA revealed a trend toward increased activity during the dark phase (*p* = 0.088) and a significant increase during the light phase (*p* = 0.0294; **Figure 1G**). Hourly analysis showed that ABA mice ran significantly less than RW controls during the food-access window (0900h: *p* = 0.0477; 1100h: *p* = 0.0176), but significantly more post-prandially (1600h: *p* = 0.0006; 1700h: *p* < 0.0001; 1800h: *p* = 0.0005) and during the light phase (0300h: *p* = 0.0043; 0400h: *p* = 0.0018; 0600h: *p* = 0.0026; 0700h: *p* = 0.0001; **Figure 1H**). These findings were confirmed when RWA was expressed relative to daily activity. ABA mice showed lower activity during the food-access window (1000h: *p* = 0.001; 1100h and 1200h: *p* < 0.0001) and at the end of the light phase (0800h: *p* = 0.0004), but greater activity post-prandially (1600h: *p* = 0.0136; 1700h: *p* = 0.0301; **Figure 1I**).

When comparing changes in total daily RWA, no significant differences were observed between the ABA and RW groups (**Figure 2A**), nor in total RWA during either the habituation or ABA periods (**Figure 2B**). However, ABA mice exhibited a significant increase in light-phase RWA compared with RW mice that was specific to the period of restricted food access (*p* = 0.0191; **Figure 2C**), whereas no significant difference between groups was found in dark-phase running activity (**Figure 2D**). This pattern was reflected in the shift observed in RWA in ABA mice towards the light phase from the commencement of ABA conditions (**Figure 2E**). Interestingly, body weight loss in ABA mice was strongly correlated with light-phase RWA (*p* = 0.0022; **Figure 2F**), but not with dark-phase running activity (**Figure 2G**), which suggests that the maladaptive hyperactivity characteristic of this model is underpinned by circadian shifts in wheel running.

**Figure 2.**
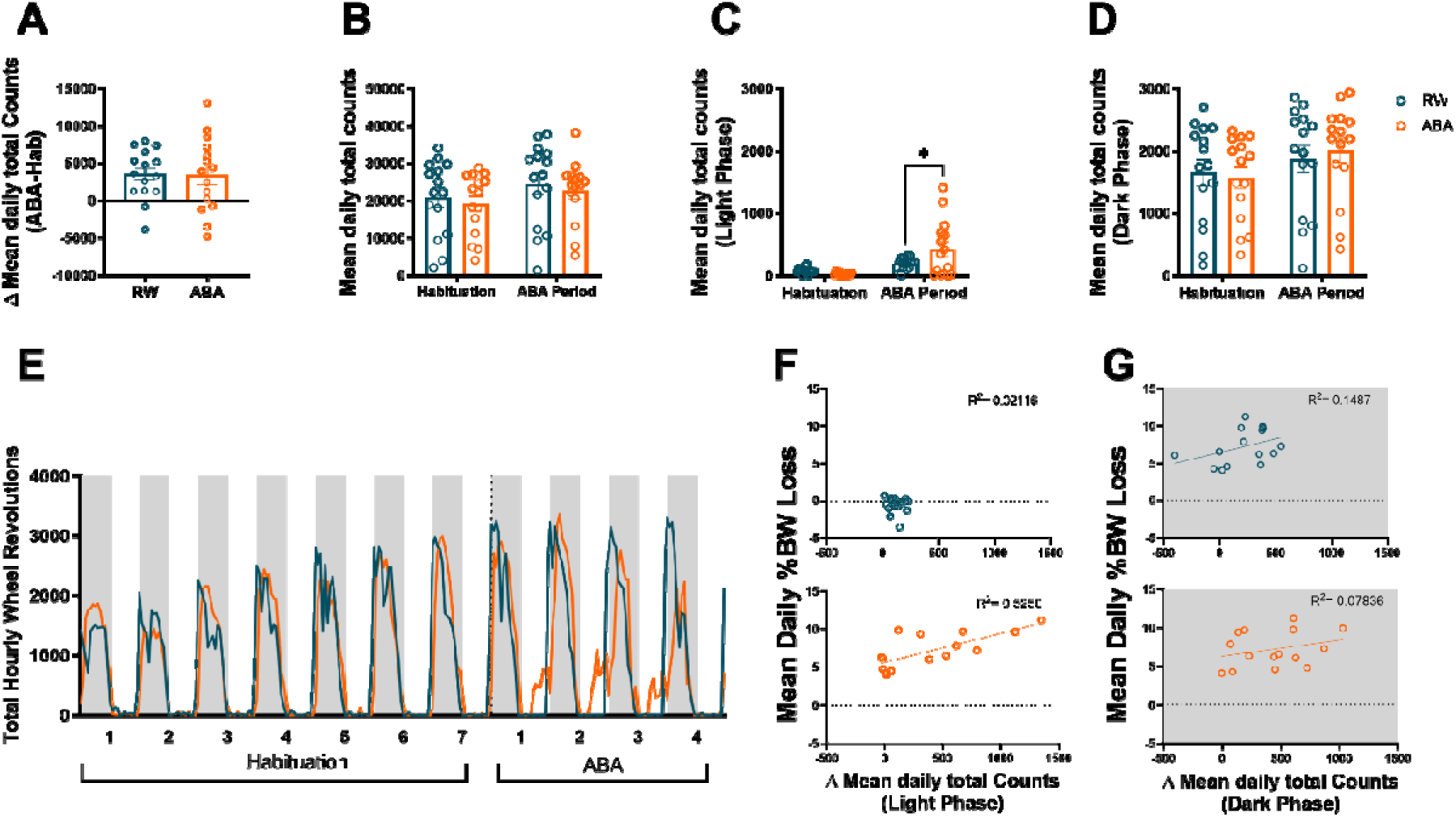
Light-phase running wheel activity is specifically elevated in ABA mice and correlates with weight loss. (A, B, D-E) No significant differences were observed between RW and ABA mice in the change of mean daily running wheel activity (RWA), overall mean daily RWA, or dark-phase RWA across both habituation and ABA periods. (C) During the light phase, ABA mice exhibited higher RWA than RW controls only during the ABA period (p = 0.0191). (F) During the light phase, RW group did not show a strong correlation between the % body weight loss during the ABA period and Δ Light-phase RWA, whereas a strong positive correlation was seen in the ABA group (Pearson’s r = 0.7251, p = 0.0022). (G) In both the RW and ABA group, no significant correlations between the % body weight loss during the ABA period and Δ dark-phase RWA were observed. Shaded areas represent the dark phase of the light cycle. ABA (Activity-Based Anorexia) and RW (running wheel) groups. Data are presented as mean ± SEM. Analyses were performed using an unpaired t-test as well as a two-way ANOVA or mixed-effects models followed by Šidák’s post hoc comparisons. Pearson correlation analyses were performed for panels F and G. *p < 0.05.

### Psilocybin treatment did not consistently alter social behaviour in ABA mice

When we evaluated the effects of PSI on sociability, we did not observe effects in ABA mice or the RW or the FR groups; however, psilocybin reduced the preference of Control mice for novel conspecific interaction when they had the option of interacting with a familiar conspecific (**Supplementary Data 1**). Using a more statistically rigorous approach of combining all conditions and drug treatments we did not observe significant effects of PSI on social behaviour (**Supplementary Figure 2**), although some interesting social outcomes emerged in SAL-treated mice, most notably an elevation in exploration of the novel over familiar conspecific mouse in ABA mice compared to all three control conditions (**Supplementary Figure 2G**). Among PSI-treated mice, those with prior access to exercise (either ABA or RW) exhibited a greater preference to interact with a novel mouse over a novel object compared to FR mice (**Supplementary Figure 2B,F,J**), suggesting an interaction between psilocybin and environmental context. Additionally, PSI-treated ABA mice exhibited a greater preference for novelty than Controls or RW mice, which was not observed in SAL-treated ABA mice (**Supplementary** Figure 2D**, H**). Motivated to understand why FR mice administered psilocybin were more interested in the novel object than the novel mouse, we correlated body weight on the day of testing with SI scores to reveal a strong positive correlation that was not observed in other conditions, or in SAL-treated FR mice (*p* = 0.0049; **Supplementary Figure 3**). This indicates that PSI increased sniffing time directed toward the novel object, particularly in mice with lower body weight, which may reflect enhanced food-seeking behaviour. Overall, since no consistent treatment effects of PSI were observed within groups, subsequent analyses focused on condition-dependent effects irrespective of drug treatment.

### Both ABA and RW mice displayed preference towards novel mouse compared to a novel object

To assess the effects of ABA exposure on sociability during the social preference trial, we evaluated locomotor activity and sociability across the total duration of the trial as well as during the exploratory phase (first 5min) and choice phase (second 5min) of the trial. Locomotion and sociability Z-scores did not differ between the four experimental groups across the full trial duration (**Figure 3A, B**). Contrary to expectations and irrespective of drug treatment, ABA mice showed an increase in sociability compared to food restricted mice (*p* = 0.0278), indicating a preference for the novel mouse over the novel object. This may be related to the exercise component of the model considering that the RW group showed a significantly higher sociability compared to both Controls (*p* = 0.0123) and FR mice (*p* = 0.0012; **Figure 3C**). No group differences in sociability Z-scores were evident during either the exploratory or choice phases (**Figure 3D, F**). However, sociability calculated from the exploratory phase indicated that ABA mice exhibited significantly greater novelty-seeking behaviour compared with controls and FR mice (*p* _Control_ = 0.0136, *p* _FR_ = 0.020; **Figure 3E**). This enhanced novelty preference persisted throughout the trial, with ABA mice exhibiting a higher sociability than FR mice during the choice phase (*p* = 0.0366). In contrast, the preference for the novel mouse observed in RW mice during the full trial was primarily driven by the choice phase, during which they showed a significantly higher sociability than both Control and FR mice (*p* _Control_ = 0.0102, *p* _FR_ = 0.0005; **Figure 3G**).

**Figure 3.**
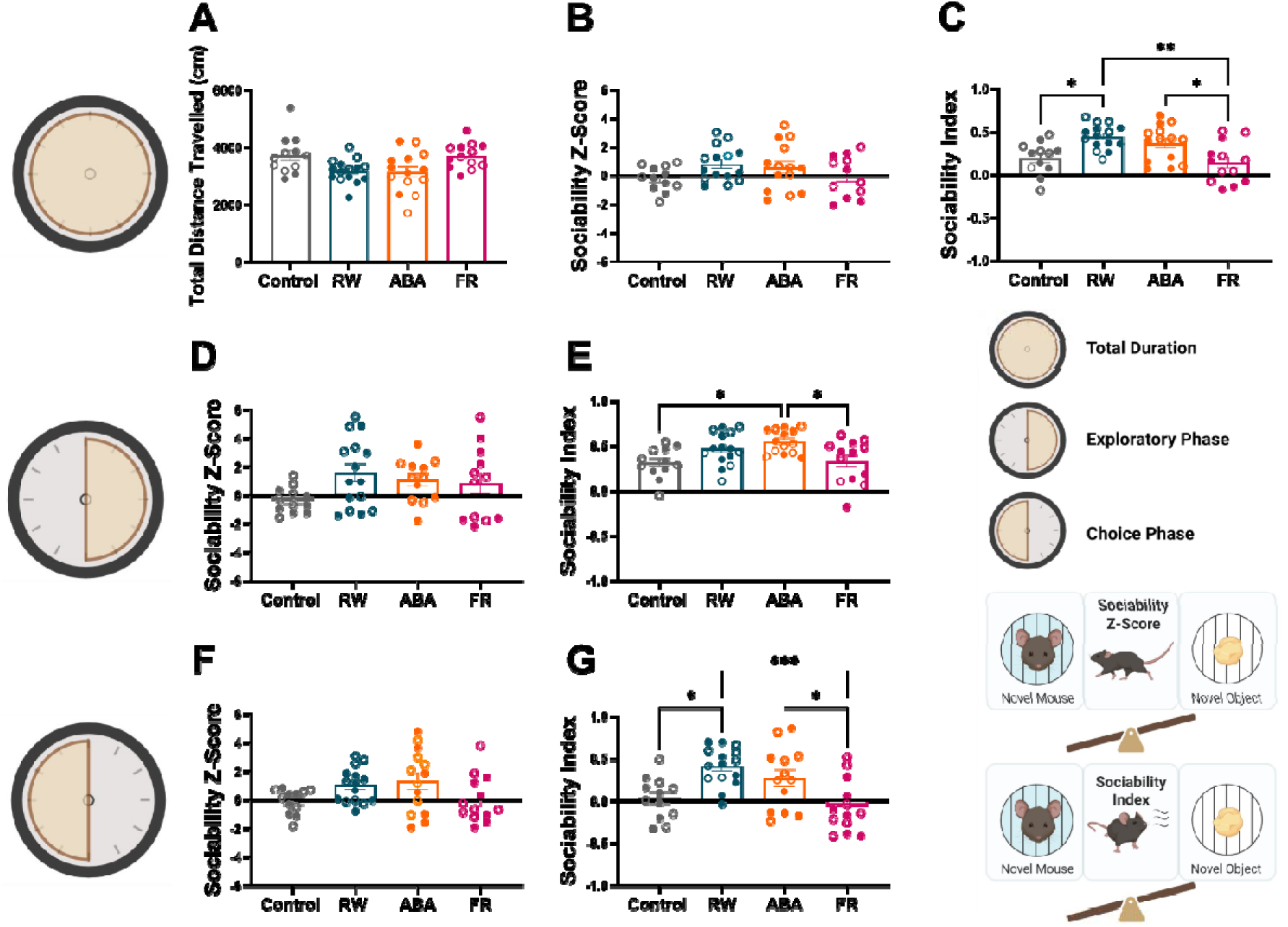
Both ABA and RW groups demonstrate elevated preference for novel social over other novel stimuli. (A) Total locomotor activity and (B) sociability Z-scores did not differ significantly between groups across the full trial duration however, (C) RW mice exhibited higher sociability than both controls (p = 0.0123) and FR mice (p = 0.0012), while ABA mice also showed higher sociability than FR mice (p = 0.0278). (D, F) No significant group differences in sociability Z-scores were observed during the exploratory phase or choice phase. (E) During the exploratory phase, ABA mice had higher sociability than controls (p = 0.0136) and FR mice (p = 0.020). (G) In the choice phase, RW mice had higher sociability than controls (p = 0.0102) and FR mice (p = 0.0005), and ABA mice had higher sociability than FR mice (p = 0.0366). ABA (Activity-Based Anorexia), FR (food-restricted), and RW (running wheel) groups. Empty symbols represent saline-treated mice; filled symbols represent psilocybin-treated mice. Sociability Z-scores were calculated from time spent and frequency of entries into the novel mouse versus novel object chambers, normalised to control values. Sociability was determined from the total time each mouse spent actively sniffing the novel mouse versus the novel object. Data are presented as mean ± SEM and were analysed by one-way ANOVA with Šidák post hoc tests. Significance thresholds: *p < 0.05; **p < 0.01; *** p < 0.001.

When considering the parameters that contribute to the sociability Z-score and sociability separately, including the duration and frequency that mice spent in either of the chambers, sniffing and interacting with the novel mouse and the novel object, the most consistent finding was an increase in novel object exploration by FR mice, again suggesting an enhanced drive for food-seeking over social interaction. In addition, all three ABA-associated groups demonstrated increased sniffing directed at the novel mouse compared to Controls (**Supplementary Data 4**).

### ABA mice exhibited a preference for novelty over familiarity during the social novelty trial

When assessing preference for novelty versus familiarity, no differences in locomotion were observed between groups across the total trial duration (**Figure 4A**). However, ABA mice showed a significantly higher preference for novelty compared to Controls, as reflected by sociability Z-scores (*p* = 0.0209; **Figure 4B**), with a trend toward significance in their SI (*p* = 0.0606). Unlike the similar trends for mice with prior exercise in the social novelty trial, ABA mice also exhibited a higher sociability compared to the RW group during this trial (*p* = 0.0059; **Figure 4C**), which highlights the distinct behavioural drivers of the two types of social preference measured by this task. During the exploration phase, ABA mice showed the highest sociability Z-scores among all groups (ABA vs Control: *p* = 0.0059; ABA vs RW: *p* = 0.0027; ABA vs FR: *p* = 0.0001; **Figure 4D**). Consistently, their sociability was significantly higher than the RW group (*p* = 0.0112) and approached significance relative to FR mice (*p* = 0.0537; **Figure 4E**). In the choice phase, no significant differences were detected between groups in either sociability Z-scores or sociability (**Figure 4F, G**). This preference of ABA mice towards novelty was reflected by elevations in all contributing parameters including time spent and frequency of entries into the novel chamber, as well as heightened sniffing and interaction with the novel mouse compared to the familiar mouse (**Supplementary Data 5A-D**).

**Figure 4.**
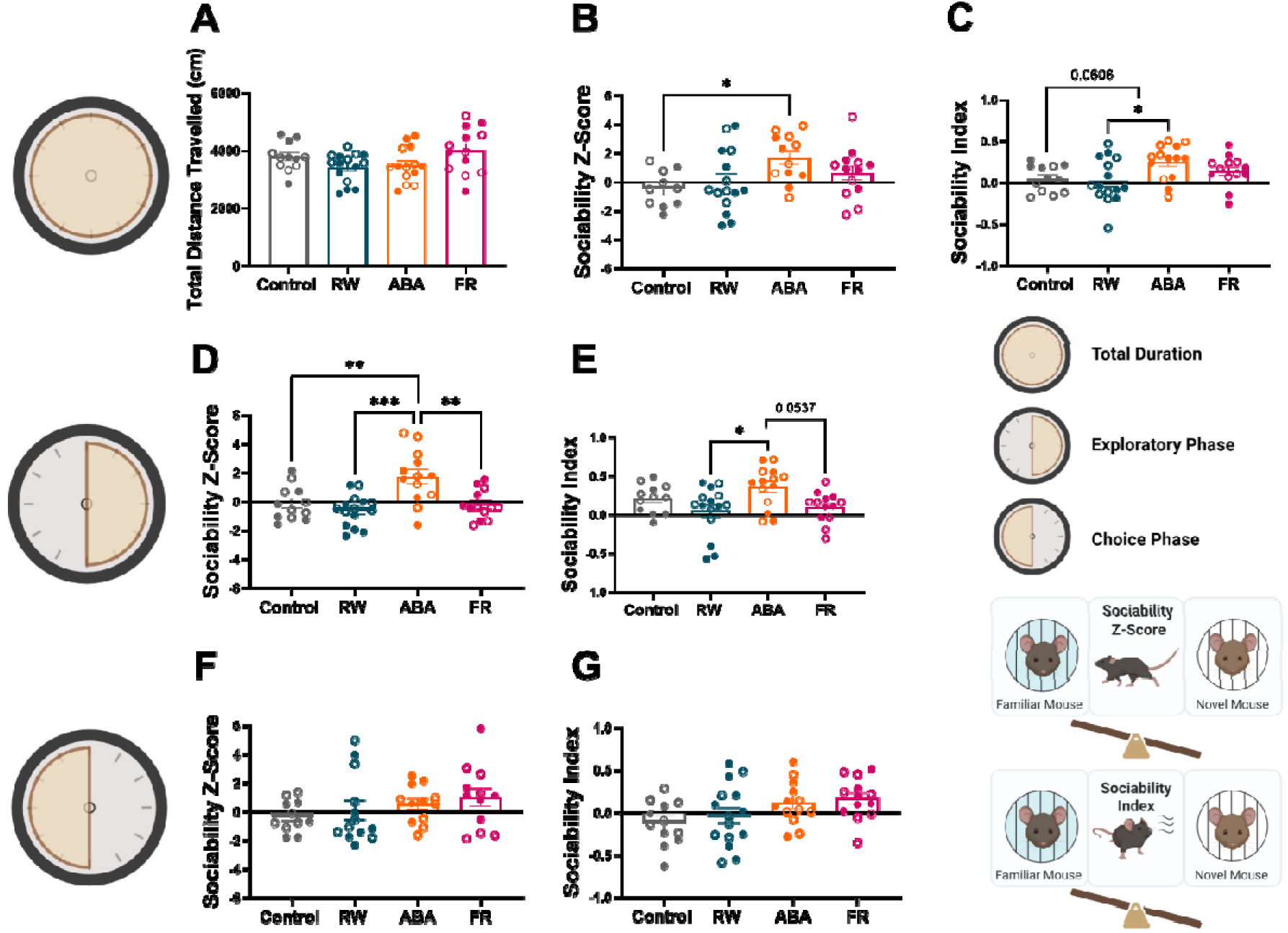
ABA mice show greater interactions with novel over familiar social stimuli, driven by behaviour in the initial exploratory phase of the task. (A) Total locomotor activity did not differ significantly between groups. (B) During the total duration of the trial, ABA mice exhibited higher Z-scores than controls (p = 0.0209) and (C) they exhibited higher sociability than RW mice (p = 0.0181) and showed a trend compared to controls (p = 0.0606). (D) ABA mice showed a significantly higher Z-score compared than controls (p = 0.0059), RW (p = 0.0027) and FR (p = 0.0001). (E) Furthermore, during the same phase, ABA mice showed a higher sociability than RW (p = 0.0112) and a trend compared to FR (p = 0.0537). (F–G) No significant group differences were observed in sociability Z-scores or sociability during the choice phase. ABA (Activity-Based Anorexia), FR (food-restricted), RW (running wheel) groups. Empty symbols represent saline-treated mice; filled symbols represent psilocybin-treated mice. Sociability Z-scores were calculated from time spent and frequency of entries into the novel versus familiar mouse chambers, normalised to control values. Sociability was determined from the total time each mouse spent actively sniffing the novel mouse versus the novel object. Data are mean ± SEM, analysed by one-way ANOVA with Šidák post hoc tests. Significance levels are indicated as: *p < 0.05; **p < 0.01; *** p < 0.001.

### Psilocybin administration elevated IL-6 levels in RW mice

When evaluating IL-6 levels across groups, exposure to different conditions (RW, ABA or FR) did not alter IL-6 concentrations in mice (**Figure 5A**) compared to Controls. However, PSI treatment significantly increased IL-6 levels in the RW group compared to SAL-treated RW mice (*p* = 0.0029), as well as compared to PSI-treated ABA (*p* = 0.011) and Control (*p* = 0.005) groups (**Figure 5B**). Moreover, IL-6 levels in the PSI-treated RW group showed a strong correlation with social interaction during the social novelty trial (*p* = 0.0166), with a similar pattern observed in PSI-treated controls (*p* = 0.0224; **Figure 5C**). Finally, when assessing behavioural influences within the ABA group, IL-6 levels were strongly and positively correlated with food intake in PSI-treated ABA mice (**Supplementary Data 6B**), suggesting a potential link between inflammatory signalling and feeding behaviour.

**Figure 5.**
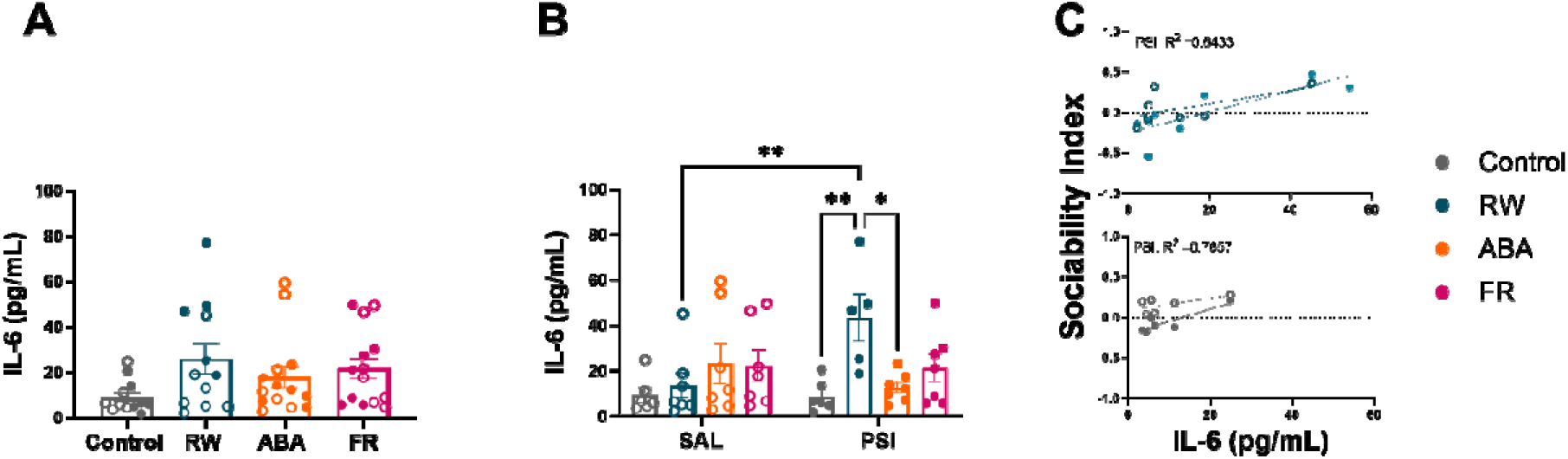
IL-6 levels are increased in RW mice administered psilocybin, which was correlated with a greater preference to interact with novel over familiar conspecifics. (A) IL-6 levels did not differ significantly between groups overall. (B) PSI-treated RW mice consistently exhibited higher IL-6 levels compared with SAL-treated RW mice (p = 0.0029), PSI-treated controls (p = 0.005) and PSI-treated ABA mice (p = 0.011). (C) A strong positive correlation was found between IL-6 levels and sociability during the social novelty trial in PSI-treated RW group (Pearson’s r = 0.8020, p = 0.0166) and in the PSI-treated controls (Pearson’s r = 0.8751, p = 0.0224). ABA (Activity-Based Anorexia), FR (food-restricted), RW (running wheel) groups, psilocybin (PSI), Saline (SAL). Empty symbols represent SAL-treated mice; filled symbols represent PSI-treated mice. Sociability was determined from the total time each mouse spent actively sniffing the novel mouse versus the novel object. Data are presented as mean ± SEM and were analysed using two-way ANOVA with Tukey’s post hoc tests and Pearson correlation was performed between the SAL and PSI-treated RW group and control. Significance levels: *p < 0.05; **p < 0.01.

## Discussion

The ABA rodent model has been widely used to investigate the core behavioural and neurobiological features of AN (Milton et al., 2018; Wable et al., 2015; Welch et al., 2021). However, whether this model in mice also replicates the social deficits associated with AN has not yet been evaluated. In parallel, clinical research on the efficacy of psilocybin in AN is ongoing, with four trials completed and two currently recruiting (ClinicalTrials.gov). To date, only one study has reported that a single dose of psilocybin is safe in females with AN (Peck et al., 2023). The present study therefore provides novel insights into how social behaviour is altered in the ABA mouse model and shows subtle effects of psilocybin on specific aspects of social behaviour in food restricted and exercising mice. By examining its effects on both behaviour and inflammation, this work contributes to a broader understanding of the mechanisms of action of psilocybin in the context of complex psychiatric conditions.

We successfully established the ABA model in female mice, as they exhibited ∼20% body weight loss along with increased postprandial and light-phase running activity, consistent with previous reports in ABA rodents (Beeler & Burghardt, 2021; Mattioni et al., 2025; Wu et al., 2014). Compared to Controls, ABA mice showed a greater preference for social novelty over familiarity, which reflected goal-directed exploratory motivation rather than random hyperactivity or anxiety, as locomotor activity did not differ across groups. These findings differ from reports in weight-restored adult female rats, which display reduced social interaction and heightened anxiety (Karth & Kinzig, 2024). Moreover, the experimental design in this study differed substantially, as rats underwent two bouts of ABA exposure with varying food availability regimens. Such discrepancies may reflect age and species differences and the influence of weight restoration on social behaviour. Recent work in adolescent mice supports this interpretation, showing that hunger-sensing agouti-related peptide (AgRP) neurons in the arcuate nucleus of the hypothalamus, which are known for being involved in regulating hunger in adults, are also activated by social isolation during adolescence (Iyilikci et al., 2025). Furthermore, AgRP neuron activity has been implicated in the mitigation of ABA symptoms, suggesting a modulatory role in the behavioural adaptations observed under food restriction (Sutton Hickey et al., 2023). Together, these findings align with our results and highlight an interplay between metabolic and social motivational systems in ABA mice.

The enhanced novelty-seeking behaviour observed following acute ABA exposure may therefore represent a homeostatic drive toward novelty as a compensatory response to restricted reward (food) or an increased motivation for social interaction, as recently reported. However, considering that this effect was absent in FR mice, an alternative explanation could be that novelty-seeking in ABA mice reflects the shared compulsive features between AN and substance use disorders (SUD) (Howard et al., 2020; López-Torrecillas, 2025) and could be considered an “addiction-prone” phenotype. There is a high prevalence of substance use and SUD in individuals with AN (Devoe et al., 2021), and in rodents, heightened novelty-seeking behaviour (in a similar manner to high impulsivity) predicts vulnerability to transition from controlled to compulsive cocaine use, even in the absence of baseline hyperactivity (Belin et al., 2011; Belin & Deroche-Gamonet, 2012; Martínez-Caballero et al., 2025). Moreover, the notion that excessive exercise exhibits addictive properties in individuals with AN (Klein et al., 2004), who also experience emotional dysregulation and novelty-seeking traits (Lozano-Madrid et al., 2020; Wonderlich et al., 2004), further strengthens this conceptual link. These behaviours share common genetic, neurobiological and theoretical frameworks with addiction models proposed for both SUD and eating disorders (Eskander et al., 2020). Acute exposure to ABA conditions during adolescence may act as an early-life stressor, heightening vulnerability to maladaptive reward processing and compulsive behaviours, including substance use. This aligns with evidence that the onset of AN often occurs during adolescence which is a critical period for the emergence of impulsivity and addictive behaviours (De Wit, 2009; Merikangas et al., 2010). Although our ABA model successfully replicated several behavioural and physiological characteristics of AN, it did not reproduce the social deficits commonly observed in patients. These differences may therefore be driven by either the psychosocial factors that shape the social deficits observed in individuals with AN or the neural adaptations that occur following sustained malnutrition. Future research should examine the impact of longer ABA exposure time on social behaviour in the mouse model, including multiple “rounds” of ABA conditions to better capture the chronic and relapsing nature of AN in humans.

To our knowledge, this is the first study to examine the influence of psilocybin on sociability in female mice exposed to ABA as well as in conditions where mice were food restricted or had access to running wheels. We observed no significant effect of psilocybin on sociability across these groups. In line with previous findings, psilocybin did not alter social preference in wild-type (WT) female mice that preferred interacting with a novel mouse over a novel object (Gattuso et al., 2025; Lu et al., 2025). Interestingly, we showed that psilocybin reduced novelty-seeking behaviour in WT (Control) females, as these mice spent equivalent time interacting with familiar and novel mice. This finding may relate to familiarity-seeking behaviour, which has been shown to be more prominent in female mice that experience a stronger reward response when engaging with familiar social partners (Xiao et al., 2025). Nonetheless, given the limited consistency of this observation and the lack of literature on the effects of psilocybin on familiarity-related behaviours, definitive conclusions cannot yet be drawn. Historically, most rodent studies investigating the effects of psilocybin on sociability have used male subjects, despite growing evidence that psychedelics may exert sex-specific effects (Shadani et al., 2024). Considering that AN predominantly affects females, this work contributes to a renewed effort to understand the mechanisms of psychedelics relevant for female-specific conditions.

Regarding the increased social exploration observed in the ABA and RW groups in our study, it is important to consider the influence of exercise on sociability. As noted previously, the detrimental effects of excessive exercise in combination with food restriction in the ABA paradigm typically require longer exposure periods to emerge (Achamrah et al., 2017; Beeler & Burghardt, 2021; Rokot et al., 2021). In the present study, both ABA and RW mice exhibited enhanced sociability, consistent with findings in female rats provided with running wheel access (Karth & Kinzig, 2024). Despite these similarities, the nature of social preference differed between exercising groups. ABA mice demonstrated sustained novelty-seeking behaviour across both the social preference and social novelty trials, whereas RW mice showed a preference for the novel mouse only during the social preference trial. This suggests that food restriction in the ABA model may amplify exploratory motivation beyond that driven by exercise alone. The pronounced novelty-seeking behaviour of ABA mice could reflect an adaptive response to food scarcity, where animals increase exploration of novel stimuli as a potential foraging strategy (Santiago et al., 2021). This effect was not evident in the FR group, likely due to their comparatively minimal body weight loss at the time of testing (approximately 5%), which may have been insufficient to elicit a similar motivational drive.

From a motivational perspective, voluntary wheel running in RW mice is inherently rewarding and activates mesolimbic dopaminergic pathways (Greenwood et al., 2011), while ABA mice experience a motivational conflict between hunger and exploration. This competing drive may enhance goal-directed novelty-seeking as a compensatory mechanism for restricted access to primary rewards such as food. These observations parallel the behavioural and motivational disturbances seen in individuals with AN, who often display heightened activity, compulsive exercise and altered reward processing (Kaye et al., 2013). In particular, increased novelty-seeking and restlessness have been associated with compulsive exercise and impaired sensitivity to natural rewards in AN (Davis & Woodside, 2002; O’Hara et al., 2016). Thus, the behavioural profile observed in ABA mice may model aspects of the reward dysregulation and compulsive tendencies characteristic of AN, highlighting the potential of this paradigm to investigate shared neurobiological mechanisms underlying maladaptive motivation across eating and addiction-related disorders.

Contrary to our expectations, ABA mice did not exhibit elevated IL-6 levels, and a single dose of psilocybin did not alter IL-6 concentrations in these mice. Our findings of unaltered IL-6 serum levels in ABA mice align with previous reports in weight-restored rats and mice (Belmonte et al., 2016; Milton et al., 2022). However, other studies have shown increased IL-6 gene expression alongside reductions in other inflammatory markers in ABA rats (Spero et al., 2024). These discrepancies may reflect differences in the duration of ABA exposure or the timing of sample collection, as IL-6 transcription and secretion follow a rapid but transient course that may not have been captured at our sampling time point. Although altered inflammatory profiles have been consistently reported in people with AN, evidence on IL-6 levels remains mixed. Some studies report reduced serum IL-6 concentrations, particularly in younger individuals (Guerrero et al., 2022; Specht et al., 2022), whereas others describe elevated IL-6 levels in adult or chronic AN patients (Dalton, Campbell, et al., 2018; Solmi et al., 2015). These contrasting findings may reflect age-related immune adaptations, disease stage or the dual pro- and anti-inflammatory roles of IL-6 (Del Giudice & Gangestad, 2018; Fuster & Walsh, 2014). Consequently, IL-6 modulation in AN may reflect a dynamic balance between inflammatory activation and compensatory regulation across the illness trajectory. In contrast to the ABA mice, psilocybin treatment in RW mice induced a significant increase in IL-6 levels. Notably, IL-6 concentrations correlated positively with SI in both PSI-treated RW and Control mice, which was not observed in SAL-treated mice in either condition. However, no such relationship between psilocybin, IL-6 and sociability was evident in either ABA or FR groups, suggesting that prior food restriction may disrupt this link.

Psychedelics, including psilocybin, act primarily through 5-HT2A receptor activation and have been shown to exert anti-inflammatory effects, partly through inhibition of TNF-α-induced inflammation and downstream mediators such as IL-6 (Flanagan & Nichols, 2018). Given that IL-6 expression is typically induced downstream of TNF-α, psilocybin’s reported inhibition of TNF-α signalling may contribute to its delayed reduction in IL-6. Thus, the absence of psilocybin-induced changes in IL-6 in our study may reflect either the acute time frame of assessment or a lack of sustained inflammatory activation in this model. The literature on psychedelics and inflammation remains limited and somewhat inconsistent with most findings derived from in vitro studies (Smedfors et al., 2022; Szabo et al., 2014). In vivo, DMT reduced systemic IL-6 levels in male rats 25 hours post-ischaemic injury (Nardai et al., 2020), and psilocybin decreased IL-6 gene expression (but not protein levels) four hours after administration in an inflammation model in male mice (Zanikov et al., 2023). In healthy humans, no acute change in IL-6 was detected following psilocybin administration, but a reduction was observed seven days post-dose, which correlated with sustained improvements in mood and social connectedness (Mason et al., 2023). Together, these findings suggest that psilocybin’s anti-inflammatory effects may emerge over time, particularly under conditions of chronic immune activation or neuroinflammatory stress, rather than during acute exposure. This delayed immunomodulation is consistent with its long-lasting psychological and therapeutic effects observed in individuals with depression and anxiety.

In summary, the present study revealed subtle and context-specific alterations of social behaviour elicited by psilocybin in female mice. These results highlight the substantial gap that remains in our understanding of the timing and mechanistic dynamics of psilocybin’s effects, particularly regarding the cascade of neuroplasticity-related events induced following 5-HTR binding. The temporal window during which psilocybin modulates synaptic plasticity, inflammation and social behaviour remains unclear, and this uncertainty is further compounded by potential sex differences in pharmacokinetics and neural processing. Male and female subjects may differ not only in how psychedelics are metabolised but also in how their neural circuits respond to serotonergic modulation. Accordingly, future research should systematically examine the effects of psilocybin and related compounds across both sexes and at multiple time points post-administration. This approach will be critical for identifying sex-specific trajectories of neuroplastic and behavioural change, especially in the context of psychedelics’ prosocial and therapeutic effects. Understanding these temporal and biological variables may clarify how psilocybin promotes enduring behavioural and affective improvements, and could help refine its translational potential for conditions such as AN.

## Acknowledgements

The authors gratefully acknowledge the USONA Institute Investigational Drug Supply Program for providing the compound used for these studies.

## References

1. Achamrah, N., Nobis, S., Goichon, A., Breton, J., Legrand, R., do Rego, J. L., do Rego, J. C., Déchelotte, P., Fetissov, S. O., Belmonte, L., & Coëffier, M. (2017). Sex differences in response to activity-based anorexia model in C57Bl/6 mice. Physiology & Behavior, 170, 1–5. 10.1016/j.physbeh.2016.12.014

2. Allen, P. J., Jimerson, D. C., Kanarek, R. B., & Kocsis, B. (2017). Impaired reversal learning in an animal model of anorexia nervosa. Physiol Behav, 179, 313–318. 10.1016/j.physbeh.2017.06.013

3. Aoki, C. (2021). Activity-based anorexia, an animal model of anorexia nervosa for investigating brain plasticity underlying the gain of resilience. Animal models of eating disorders, 267–296.

4. Association, A. P. (2013). Diagnostic and statistical manual of mental disorders: DSM-5. American psychiatric association.

5. Beeler, J. A., & Burghardt, N. S. (2021). Activity-based Anorexia for Modeling Vulnerability and Resilience in Mice. Bio Protoc, 11(9), e4009. 10.21769/BioProtoc.4009

6. Belin, D., Berson, N., Balado, E., Piazza, P. V., & Deroche-Gamonet, V. (2011). High-novelty-preference rats are predisposed to compulsive cocaine self-administration. Neuropsychopharmacology, 36(3), 569–579. 10.1038/npp.2010.188

7. Belin, D., & Deroche-Gamonet, V. (2012). Responses to novelty and vulnerability to cocaine addiction: contribution of a multi-symptomatic animal model. Cold Spring Harb Perspect Med, 2(11). 10.1101/cshperspect.a011940

8. Belmonte, L., Achamrah, N., Nobis, S., Guérin, C., Riou, G., Bôle-Feysot, C., Boyer, O., Richard, V., Rego, J. C., Déchelotte, P., Goichon, A., & Coëffier, M. (2016). A role for intestinal TLR4- driven inflammatory response during activity-based anorexia. Sci Rep, 6, 35813. 10.1038/srep35813

9. Bevilacqua, A., Santini, F., La Porta, D., & Cimino, S. (2024). Association of serotonin receptor gene polymorphisms with anorexia nervosa: a systematic review and meta-analysis. Eat Weight Disord, 29(1), 31. 10.1007/s40519-024-01659-3

10. Bhatt, K. V., & Weissman, C. R. (2024). The effect of psilocybin on empathy and prosocial behavior: a proposed mechanism for enduring antidepressant effects. npj Mental Health Research, 3(1), 7. 10.1038/s44184-023-00053-8

11. Bora, E., & Köse, S. (2016). Meta-analysis of theory of mind in anorexia nervosa and bulimia nervosa: A specific İmpairment of cognitive perspective taking in anorexia nervosa? International Journal of Eating Disorders, 49(8), 739–740. 10.1002/eat.22572

12. Cadirci, E., Halici, Z., Bayir, Y., Albayrak, A., Karakus, E., Polat, B., Unal, D., Atamanalp, S. S., Aksak, S., & Gundogdu, C. (2013). Peripheral 5-HT7 receptors as a new target for prevention of lung injury and mortality in septic rats. Immunobiology, 218(10), 1271–1283.

13. Capuron, L., & Miller, A. H. (2011). Immune system to brain signaling: Neuropsychopharmacological implications. Pharmacology & Therapeutics, 130(2), 226–238. 10.1016/j.pharmthera.2011.01.014

14. Carhart-Harris, R. L., Bolstridge, M., Day, C. M. J., Rucker, J., Watts, R., Erritzoe, D. E., Kaelen, M., Giribaldi, B., Bloomfield, M., Pilling, S., Rickard, J. A., Forbes, B., Feilding, A., Taylor, D., Curran, H. V., & Nutt, D. J. (2018). Psilocybin with psychological support for treatment-resistant depression: six-month follow-up. Psychopharmacology, 235(2), 399–408. 10.1007/s00213-017-4771-x

15. Carhart-Harris, R. L., Bolstridge, M., Rucker, J., Day, C. M. J., Erritzoe, D., Kaelen, M., Bloomfield, M., Rickard, J. A., Forbes, B., Feilding, A., Taylor, D., Pilling, S., Curran, V. H., & Nutt, D. J. (2016). Psilocybin with psychological support for treatment-resistant depression: an open-label feasibility study. The Lancet Psychiatry, 3(7), 619–627. 10.1016/S2215-0366(16)30065-7

16. Conn, K., Milton, L., Huang, K., Munguba, H., Ruuska, J., Lemus, M., Greaves, E., Homman-Ludiye, J., Oldfield, B., & Foldi, C. (2024). Psilocybin prevents activity-based anorexia in female rats by enhancing cognitive flexibility: contributions from 5-HT1A and 5-HT2A receptor mechanisms. bioRxiv, 2023.2012.2012.571374. 10.1101/2023.12.12.571374

17. Costa, L. H. A., Santos, B. M., & Branco, L. G. S. (2020). Can selective serotonin reuptake inhibitors have a neuroprotective effect during COVID-19? European Journal of Pharmacology, 889, 173629. 10.1016/j.ejphar.2020.173629

18. Dakic, V., Minardi Nascimento, J., Costa Sartore, R., Maciel, R. M., de Araujo, D. B., Ribeiro, S., Martins-de-Souza, D., & Rehen, S. K. (2017). Short term changes in the proteome of human cerebral organoids induced by 5-MeO-DMT. Sci Rep, 7(1), 12863. 10.1038/s41598-017-12779-5

19. Dalton, B., Bartholdy, S., Robinson, L., Solmi, M., Ibrahim, M. A., Breen, G., Schmidt, U., & Himmerich, H. (2018). A meta-analysis of cytokine concentrations in eating disorders. Journal of psychiatric research, 103, 252–264.

20. Dalton, B., Campbell, I. C., Chung, R., Breen, G., Schmidt, U., & Himmerich, H. (2018). Inflammatory Markers in Anorexia Nervosa: An Exploratory Study. Nutrients, 10(11), 1573. https://www.mdpi.com/2072-6643/10/11/1573

21. Davis, A. K., Barrett, F. S., May, D. G., Cosimano, M. P., Sepeda, N. D., Johnson, M. W., Finan, P. H., & Griffiths, R. R. (2021). Effects of Psilocybin-Assisted Therapy on Major Depressive Disorder: A Randomized Clinical Trial. JAMA Psychiatry, 78(5), 481–489. 10.1001/jamapsychiatry.2020.3285

22. Davis, C., & Woodside, D. B. (2002). Sensitivity to the rewarding effects of food and exercise in the eating disorders. Compr Psychiatry, 43(3), 189–194. 10.1053/comp.2002.32356

23. De Gregorio, D., Aguilar-Valles, A., Preller, K. H., Heifets, B. D., Hibicke, M., Mitchell, J., & Gobbi, G. (2021). Hallucinogens in Mental Health: Preclinical and Clinical Studies on LSD, Psilocybin, MDMA, and Ketamine. J Neurosci, 41(5), 891–900. 10.1523/jneurosci.1659-20.2020

24. De Wit, H. (2009). Impulsivity as a determinant and consequence of drug use: a review of underlying processes. Addiction biology, 14(1), 22–31.

25. DeJong, H., Van den Eynde, F., Broadbent, H., Kenyon, M. D., Lavender, A., Startup, H., & Schmidt, U. (2013). Social cognition in bulimia nervosa: a systematic review. European Psychiatry, 28(1), 1–6.

26. Del Giudice, M., & Gangestad, S. W. (2018). Rethinking IL-6 and CRP: Why they are more than inflammatory biomarkers, and why it matters. Brain, Behavior, and Immunity, 70, 61–75. 10.1016/j.bbi.2018.02.013

27. Devoe, D. J., Dimitropoulos, G., Anderson, A., Bahji, A., Flanagan, J., Soumbasis, A., Patten, S. B., Lange, T., & Paslakis, G. (2021). The prevalence of substance use disorders and substance use in anorexia nervosa: a systematic review and meta-analysis. J Eat Disord, 9(1), 161. 10.1186/s40337-021-00516-3

28. Dong, Y., Goodwin-Groen, S., Ma, J., Kim, E., Giudice, S. D., Santos, M., & Aoki, C. (2025). Mechanisms underlying sustained resilience against anorexia nervosa from sub-anesthetic ketamine: a review and new research based on electron microscopic analyses of synapses using a mouse model. Physiology & Behavior, 114956. 10.1016/j.physbeh.2025.114956

29. Eskander, N., Chakrapani, S., & Ghani, M. R. (2020). The Risk of Substance Use Among Adolescents and Adults With Eating Disorders. Cureus, 12(9), e10309. 10.7759/cureus.10309

30. Flanagan, T. W., & Nichols, C. D. (2018). Psychedelics as anti-inflammatory agents. International Review of Psychiatry, 30(4), 363–375. 10.1080/09540261.2018.1481827

31. Fuster, J. J., & Walsh, K. (2014). The Good, the Bad, and the Ugly of interleukin-6 signaling. The EMBO Journal, 33(13), 1425–1427. 10.15252/embj.201488856

32. Gattuso, J. J., Kong, G., Bezcioglu, B., Lu, D., Ekwudo, M. N., Wilson, C., Gubert, C., Hannan, A. J., & Renoir, T. (2025). Chronic psilocybin administration increases sociability and alters the gut microbiome in male wild-type mice but not in a preclinical model of obsessive-compulsive disorder. Neuropharmacology, 279, 110648. 10.1016/j.neuropharm.2025.110648

33. Gibson, D., & Mehler, P. S. (2019). Anorexia Nervosa and the Immune System—A Narrative Review. Journal of Clinical Medicine, 8(11), 1915. https://www.mdpi.com/2077-0383/8/11/1915

34. Godart, N. T., Flament, M. F., Perdereau, F., & Jeammet, P. (2002). Comorbidity between eating disorders and anxiety disorders: A review. International Journal of Eating Disorders, 32(3), 253–270. 10.1002/eat.10096

35. Goodwin, G. M., Aaronson, S. T., Alvarez, O., Atli, M., Bennett, J. C., Croal, M., DeBattista, C., Dunlop, B. W., Feifel, D., Hellerstein, D. J., Husain, M. I., Kelly, J. R., Lennard-Jones, M. R., Licht, R. W., Marwood, L., Mistry, S., Páleníček, T., Redjep, O., Repantis, D., … Malievskaia, E. (2023). Single-dose psilocybin for a treatment-resistant episode of major depression: Impact on patient-reported depression severity, anxiety, function, and quality of life. Journal of Affective Disorders, 327, 120–127. 10.1016/j.jad.2023.01.108

36. Greenwood, B. N., Foley, T. E., Le, T. V., Strong, P. V., Loughridge, A. B., Day, H. E., & Fleshner, M. (2011). Long-term voluntary wheel running is rewarding and produces plasticity in the mesolimbic reward pathway. Behav Brain Res, 217(2), 354–362. 10.1016/j.bbr.2010.11.005

37. Griffiths, R. R., Johnson, M. W., Carducci, M. A., Umbricht, A., Richards, W. A., Richards, B. D., Cosimano, M. P., & Klinedinst, M. A. (2016). Psilocybin produces substantial and sustained decreases in depression and anxiety in patients with life-threatening cancer: A randomized double-blind trial. J Psychopharmacol, 30(12), 1181–1197. 10.1177/0269881116675513

38. Grob, C. S., Danforth, A. L., Chopra, G. S., Hagerty, M., McKay, C. R., Halberstadt, A. L., & Greer, G. R. (2011). Pilot study of psilocybin treatment for anxiety in patients with advanced-stage cancer. Arch Gen Psychiatry, 68(1), 71–78. 10.1001/archgenpsychiatry.2010.116

39. Gruol, D. L. (2015). IL-6 regulation of synaptic function in the CNS. Neuropharmacology, 96(Pt A), 42–54. 10.1016/j.neuropharm.2014.10.023

40. Guerrero, F. R., Gómez, J. G., Gonzalez, P. B., García, J. G., Miguel, A. B., Giraldo, G. C., García- Unzueta, M. T., & Del Barrio, A. G. (2022). Low levels of proinflammatory cytokines in a transdiagnostic sample of young male and female early onset eating disorders without any previous treatment: a case control study. Psychiatry research, 310, 114449.

41. Guilloux, J. P., Seney, M., Edgar, N., & Sibille, E. (2011). Integrated behavioral z-scoring increases the sensitivity and reliability of behavioral phenotyping in mice: relevance to emotionality and sex. J Neurosci Methods, 197(1), 21–31. 10.1016/j.jneumeth.2011.01.019

42. Haroon, E., Raison, C. L., & Miller, A. H. (2012). Psychoneuroimmunology meets neuropsychopharmacology: translational implications of the impact of inflammation on behavior. Neuropsychopharmacology, 37(1), 137–162. 10.1038/npp.2011.205

43. Harrison, A., Mountford, V. A., & Tchanturia, K. (2014). Social anhedonia and work and social functioning in the acute and recovered phases of eating disorders. Psychiatry research, 218(1-2), 187–194.

44. Herr, N., Bode, C., & Duerschmied, D. (2017). The effects of serotonin in immune cells. Frontiers in cardiovascular medicine, 4, 48.

45. Hole, C., Dhamsania, A., Brown, C., & Ryznar, R. (2025). Immune Dysregulation in Depression and Anxiety: A Review of the Immune Response in Disease and Treatment. Cells, 14(8), 607. https://www.mdpi.com/2073-4409/14/8/607

46. House, R. V., Thomas, P. T., & Bhargava, H. N. (1994). Immunological consequences of in vitro exposure to lysergic acid diethylamide (LSD). Immunopharmacol Immunotoxicol, 16(1), 23–40. 10.3109/08923979409029898

47. Howard, M., Gregertsen, E. C., Hindocha, C., & Serpell, L. (2020). Impulsivity and compulsivity in anorexia and bulimia nervosa: A systematic review. Psychiatry Res, 293, 113354. 10.1016/j.psychres.2020.113354

48. Huang, K., Milton, L. K., Dempsey, H., Power, S. J., Conn, K.-A., Andrews, Z. B., & Foldi, C. J. (2023). Rapid, automated, and experimenter-free touchscreen testing reveals reciprocal interactions between cognitive flexibility and activity-based anorexia in female rats. eLife, 12, e84961. 10.7554/eLife.84961

49. Iyilikci, O., Kim, L., Zimmer, M. R., Bober, J., Li, Y., Pelts, M., Santana, G. M., & Dietrich, M. O. (2025). Age-specific regulation of sociability by hypothalamic Agrp neurons. Current Biology, 35(18), 4522–4536.e4526. 10.1016/j.cub.2025.08.014

50. Jungwirth, J., von Rotz, R., Dziobek, I., Vollenweider, F. X., & Preller, K. H. (2024). Psilocybin increases emotional empathy in patients with major depression. Molecular Psychiatry. 10.1038/s41380-024-02875-0

51. Karth, M., & Kinzig, K. P. (2024). Adolescent activity-based anorexia has a substantial and prolonged impact on social behavior in young adult female rats. Physiology & Behavior, 279, 114528. 10.1016/j.physbeh.2024.114528

52. Kaye, W. (2008). Neurobiology of anorexia and bulimia nervosa. Physiol Behav, 94(1), 121–135. 10.1016/j.physbeh.2007.11.037

53. Kaye, W. H., Wierenga, C. E., Bailer, U. F., Simmons, A. N., & Bischoff-Grethe, A. (2013). Nothing tastes as good as skinny feels: the neurobiology of anorexia nervosa. Trends Neurosci, 36(2), 110–120. 10.1016/j.tins.2013.01.003

54. Kelley, D. P., Venable, K., Destouni, A., Billac, G., Ebenezer, P., Stadler, K., Nichols, C., Barker, S., & Francis, J. (2022). Pharmahuasca and DMT Rescue ROS Production and Differentially Expressed Genes Observed after Predator and Psychosocial Stress: Relevance to Human PTSD. ACS Chem Neurosci, 13(2), 257–274. 10.1021/acschemneuro.1c00660

55. Kiser, D., Steemers, B., Branchi, I., & Homberg, J. R. (2012). The reciprocal interaction between serotonin and social behaviour. Neurosci Biobehav Rev, 36(2), 786–798. 10.1016/j.neubiorev.2011.12.009

56. Klein, D. A., Bennett, A. S., Schebendach, J., Foltin, R. W., Devlin, M. J., & Walsh, B. T. (2004). Exercise “Addiction” in Anorexia Nervosa: Model Development and Pilot Data. CNS Spectrums, 9(7), 531–537. 10.1017/S1092852900009627

57. Köhler, C. A., Freitas, T. H., Maes, M., de Andrade, N. Q., Liu, C. S., Fernandes, B. S., Stubbs, B., Solmi, M., Veronese, N., Herrmann, N., Raison, C. L., Miller, B. J., Lanctôt, K. L., & Carvalho, A. F. (2017). Peripheral cytokine and chemokine alterations in depression: a meta-analysis of 82 studies. Acta Psychiatrica Scandinavica, 135(5), 373–387. 10.1111/acps.12698

58. Kraus, C., Castrén, E., Kasper, S., & Lanzenberger, R. (2017). Serotonin and neuroplasticity - Links between molecular, functional and structural pathophysiology in depression. Neurosci Biobehav Rev, 77, 317–326. 10.1016/j.neubiorev.2017.03.007

59. Lima Giacobbo, B., Doorduin, J., Klein, H. C., Dierckx, R. A., Bromberg, E., & de Vries, E. F. (2019). Brain-derived neurotrophic factor in brain disorders: focus on neuroinflammation. Molecular Neurobiology, 56, 3295–3312.

60. López-Torrecillas, F. (2025). Editorial: Impulsivity and compulsivity related to substance use disorders. Front Psychiatry, 16, 1599890. 10.3389/fpsyt.2025.1599890

61. Lozano-Madrid, M., Clark Bryan, D., Granero, R., Sánchez, I., Riesco, N., Mallorquí-Bagué, N., Jiménez-Murcia, S., Treasure, J., & Fernández-Aranda, F. (2020). Impulsivity, Emotional Dysregulation and Executive Function Deficits Could Be Associated with Alcohol and Drug Abuse in Eating Disorders. Journal of Clinical Medicine, 9(6), 1936. https://www.mdpi.com/2077-0383/9/6/1936

62. Lu, O. D., White, K., Raymond, K., Liu, C., Klein, A. S., Green, N., Vaillancourt, S., Gallagher, A., Shindy, L., Li, A., Wallquist, K., Li, R., Zou, M., Casey, A. B., Cameron, L. P., Pomrenze, M. B., Sohal, V., Kheirbek, M. A., Gomez, A. M., Malenka, R. (2025). A multi-institutional investigation of psilocybin’s effects on mouse behavior. bioRxiv, 2025.2004.2008.647810. 10.1101/2025.04.08.647810

63. Markopoulos, A., Inserra, A., De Gregorio, D., & Gobbi, G. (2021). Lysergic acid diethylamide (LSD) promotes social behaviour through 5-HT2A and ampa in the medial prefrontal cortex (MPFC). European Psychiatry, 64(S1), S416–S417. 10.1192/j.eurpsy.2021.1112

64. Martínez-Caballero, M., Calpe-López, C., García-Pardo, M. P., Arenas, M. C., de la Rubia Ortí, J. E., Benlloch, M., Manzanedo, C., & Aguilar, M. A. (2025). Enhanced novelty-seeking after early adolescent exposure to vicarious social defeat predicts the vulnerability of female mice to cocaine reward. Pharmacol Biochem Behav, 253, 174039. 10.1016/j.pbb.2025.174039

65. Mason, N. L., Szabo, A., Kuypers, K. P. C., Mallaroni, P. A., de la Torre Fornell, R., Reckweg, J. T., Tse, D. H. Y., Hutten, N. R. P. W., Feilding, A., & Ramaekers, J. G. (2023). Psilocybin induces acute and persisting alterations in immune status in healthy volunteers: An experimental, placebo-controlled study. Brain, Behavior, and Immunity, 114, 299–310. 10.1016/j.bbi.2023.09.004

66. Mattioni, J., Duriez, P., Aïdat, S., Lebrun, N., Bohlooly-Y, M., Gorwood, P., Viltart, O., & Tolle, V. (2025). Altered circadian pattern of activity in a chronic activity-based anorexia nervosa-like female mouse model deficient for GHSR. Psychoneuroendocrinology, 177, 107453. 10.1016/j.psyneuen.2025.107453

67. Merikangas, K. R., He, J. P., Burstein, M., Swanson, S. A., Avenevoli, S., Cui, L., Benjet, C., Georgiades, K., & Swendsen, J. (2010). Lifetime prevalence of mental disorders in U.S. adolescents: results from the National Comorbidity Survey Replication--Adolescent Supplement (NCS-A). J Am Acad Child Adolesc Psychiatry, 49(10), 980–989. 10.1016/j.jaac.2010.05.017

68. Milton, L. K., Oldfield, B. J., & Foldi, C. J. (2018). Evaluating anhedonia in the activity-based anorexia (ABA) rat model. Physiology & Behavior, 194, 324–332. 10.1016/j.physbeh.2018.06.023

69. Milton, L. K., Patton, T., O’Keeffe, M., Oldfield, B. J., & Foldi, C. J. (2022). In pursuit of biomarkers for predicting susceptibility to activity-based anorexia in adolescent female rats. International Journal of Eating Disorders, 55(5), 664–677. 10.1002/eat.23705

70. Morris, R., Bramham, J., Smith, E., & Tchanturia, K. (2014). Empathy and social functioning in anorexia nervosa before and after recovery. Cognitive neuropsychiatry, 19(1), 47–57.

71. Moy, S. S., Nadler, J. J., Perez, A., Barbaro, R. P., Johns, J. M., Magnuson, T. R., Piven, J., & Crawley, J. N. (2004). Sociability and preference for social novelty in five inbred strains: an approach to assess autistic-like behavior in mice. Genes Brain Behav, 3(5), 287–302. 10.1111/j.1601-1848.2004.00076.x

72. Nardai, S., László, M., Szabó, A., Alpár, A., Hanics, J., Zahola, P., Merkely, B., Frecska, E., & Nagy, Z. (2020). N,N-dimethyltryptamine reduces infarct size and improves functional recovery following transient focal brain ischemia in rats. Exp Neurol, 327, 113245. 10.1016/j.expneurol.2020.113245

73. Nardou, R., Sawyer, E., Song, Y. J., Wilkinson, M., Padovan-Hernandez, Y., de Deus, J. L., Wright, N., Lama, C., Faltin, S., Goff, L. A., Stein-O’Brien, G. L., & Dölen, G. (2023). Psychedelics reopen the social reward learning critical period. Nature, 618(7966), 790–798. 10.1038/s41586-023-06204-3

74. Nau, F., Jr., Yu, B., Martin, D., & Nichols, C. D. (2013). Serotonin 5-HT2A receptor activation blocks TNF-α mediated inflammation in vivo. PLoS One, 8(10), e75426. 10.1371/journal.pone.0075426

75. Nguyen, M., Allison, S., Looi, J. C., & Bastiampillai, T. (2022). Increasing hospital admission rates for anorexia nervosa amongst young women in Australia from 1998 to 2018. Australasian Psychiatry, 30(4), 462–471. 10.1177/10398562221077890

76. O’Hara, C. B., Keyes, A., Renwick, B., Leyton, M., Campbell, I. C., & Schmidt, U. (2016). The Effects of Acute Dopamine Precursor Depletion on the Reinforcing Value of Exercise in Anorexia Nervosa. PLoS One, 11(1), e0145894. 10.1371/journal.pone.0145894

77. Peck, S. K., Shao, S., Gruen, T., Yang, K., Babakanian, A., Trim, J., Finn, D. M., & Kaye, W. H. (2023). Psilocybin therapy for females with anorexia nervosa: a phase 1, open-label feasibility study. Nat Med, 29(8), 1947–1953. 10.1038/s41591-023-02455-9

78. Quintero-Villegas, A., & Valdés-Ferrer, S. I. (2020). Role of 5-HT7 receptors in the immune system in health and disease. Molecular Medicine, 26(1), 2.

79. Ravi, M., Miller, A. H., & Michopoulos, V. (2021). The immunology of stress and the impact of inflammation on the brain and behaviour. BJPsych Advances, 27(3), 158–165. 10.1192/bja.2020.82

80. Renna, M. E., O’Toole, M. S., Spaeth, P. E., Lekander, M., & Mennin, D. S. (2018). The association between anxiety, traumatic stress, and obsessive–compulsive disorders and chronic inflammation: A systematic review and meta-analysis. Depression and Anxiety, 35(11), 1081–1094. 10.1002/da.22790

81. Riddle, D. B., Guzick, A., Minhajuddin, A., Smárason, O., Armstrong, G. M., Slater, H., Mayes, T. L., Goodman, L. C., Baughn, D. L., Martin, S. L., Wakefield, S. M., Blader, J., Brown, R., Tonarelli, S., Goodman, W. K., Trivedi, M. H., & Storch, E. A. (2023). Obsessive-compulsive disorder in youth and young adults with depression: Clinical characteristics of comorbid presentations. Journal of Obsessive-Compulsive and Related Disorders, 38, 100820. 10.1016/j.jocrd.2023.100820

82. Rokot, N. T., Ataka, K., Iwai, H., Suzuki, H., Tachibe, H., Kairupan, T. S., Cheng, K.-C., Amitani, H., Inui, A., & Asakawa, A. (2021). Antagonism for NPY signaling reverses cognitive behavior defects induced by activity-based anorexia in mice. Psychoneuroendocrinology, 126, 105133. 10.1016/j.psyneuen.2021.105133

83. Ross, S., Bossis, A., Guss, J., Agin-Liebes, G., Malone, T., Cohen, B., Mennenga, S. E., Belser, A., Kalliontzi, K., Babb, J., Su, Z., Corby, P., & Schmidt, B. L. (2016). Rapid and sustained symptom reduction following psilocybin treatment for anxiety and depression in patients with life-threatening cancer: a randomized controlled trial. J Psychopharmacol, 30(12), 1165–1180. 10.1177/0269881116675512

84. Santiago, A. N., Makowicz, E. A., Du, M., & Aoki, C. (2021). Food Restriction Engages Prefrontal Corticostriatal Cells and Local Microcircuitry to Drive the Decision to Run versus Conserve Energy. Cereb Cortex, 31(6), 2868–2885. 10.1093/cercor/bhaa394

85. Saris, I. M. J., Aghajani, M., van der Werff, S. J. A., van der Wee, N. J. A., & Penninx, B. (2017). Social functioning in patients with depressive and anxiety disorders. Acta Psychiatr Scand, 136(4), 352–361. 10.1111/acps.12774

86. Singer, T. (2006). The neuronal basis and ontogeny of empathy and mind reading: Review of literature and implications for future research. Neuroscience & Biobehavioral Reviews, 30(6), 855–863. 10.1016/j.neubiorev.2006.06.011

87. Smedfors, G., Glotfelty, E., Kalani, N., Papatziamos Hjelle, C., Horntvedt, O., Wellfelt, K., Brodin, A., von Kieseritzky, F., Olson, L., & Karlsson, T. (2022). Psilocybin Combines Rapid Synaptogenic And Anti-Inflammatory Effects In Vitro. 10.21203/rs.3.rs-1321542/v1

88. Solmi, M., Veronese, N., Favaro, A., Santonastaso, P., Manzato, E., Sergi, G., & Correll, C. U. (2015). Inflammatory cytokines and anorexia nervosa: A meta-analysis of cross-sectional and longitudinal studies. Psychoneuroendocrinology, 51, 237–252. 10.1016/j.psyneuen.2014.09.031

89. Songtachalert, T., Roomruangwong, C., Carvalho, A. F., Bourin, M., & Maes, M. (2018). Anxiety Disorders: Sex Differences in Serotonin and Tryptophan Metabolism. Curr Top Med Chem, 18(19), 1704–1715. 10.2174/1568026618666181115093136

90. Specht, H. E., Mannig, N., Belheouane, M., Andreani, N. A., Tenbrock, K., Biemann, R., Borucki, K., Dahmen, B., Dempfle, A., & Baines, J. F. (2022). Lower serum levels of IL-1β and IL-6 cytokines in adolescents with anorexia nervosa and their association with gut microbiota in a longitudinal study. Frontiers in psychiatry, 13, 920665.

91. Spero, V., Scherma, M., D’Amelio, S., Collu, R., Dedoni, S., Camoglio, C., Siddi, C., Fratta, W., Molteni, R., & Fadda, P. (2024). Activity-based anorexia (ABA) model: Effects on brain neuroinflammation, redox balance and neuroplasticity during the acute phase. Neurochemistry International, 180, 105842. 10.1016/j.neuint.2024.105842

92. Sutton Hickey, A. K., Duane, S. C., Mickelsen, L. E., Karolczak, E. O., Shamma, A. M., Skillings, A., Li, C., & Krashes, M. J. (2023). AgRP neurons coordinate the mitigation of activity-based anorexia. Mol Psychiatry, 28(4), 1622–1635. 10.1038/s41380-022-01932-w

93. Szabo, A., Kovacs, A., Frecska, E., & Rajnavolgyi, E. (2014). Psychedelic N,N-dimethyltryptamine and 5-methoxy-N,N-dimethyltryptamine modulate innate and adaptive inflammatory responses through the sigma-1 receptor of human monocyte-derived dendritic cells. PLoS One, 9(8), e106533. 10.1371/journal.pone.0106533

94. Tauro, J. L., Wearne, T. A., Belevski, B., Filipčíková, M., & Francis, H. M. (2022). Social cognition in female adults with Anorexia Nervosa: A systematic review. Neuroscience & Biobehavioral Reviews, 132, 197–210. 10.1016/j.neubiorev.2021.11.035

95. Troop, N. A., & Bifulco, A. (2002). Childhood social arena and cognitive sets in eating disorders. British Journal of Clinical Psychology, 41(2), 205–211.

96. Wable, G. S., Min, J. Y., Chen, Y. W., & Aoki, C. (2015). Anxiety is correlated with running in adolescent female mice undergoing activity-based anorexia. Behav Neurosci, 129(2), 170–182. 10.1037/bne0000040

97. Weiss, F., Magnesa, A., Gambini, M., Gurrieri, R., Annuzzi, E., Elefante, C., Perugi, G., & Marazziti, D. (2025). Psychedelic-Induced Neural Plasticity: A Comprehensive Review and a Discussion of Clinical Implications. Brain Sciences, 15(2), 117. https://www.mdpi.com/2076-3425/15/2/117

98. Welch, A. C., Zhang, J., Lyu, J., McMurray, M. S., Javitch, J. A., Kellendonk, C., & Dulawa, S. C. (2021). Dopamine D2 receptor overexpression in the nucleus accumbens core induces robust weight loss during scheduled fasting selectively in female mice. Mol Psychiatry, 26(8), 3765–3777. 10.1038/s41380-019-0633-8

99. Whitten, W. K., Bronson, F. H., & Greenstein, J. A. (1968). Estrus-Inducing Pheromone of Male Mice: Transport by Movement of Air. Science, 161(3841), 584–585. doi:10.1126/science.161.3841.584

100. Wonderlich, S. A., Connolly, K. M., & Stice, E. (2004). Impulsivity as a risk factor for eating disorder behavior: Assessment implications with adolescents. International Journal of Eating Disorders, 36(2), 172–182. 10.1002/eat.20033

101. Wu, H., van Kuyck, K., Tambuyzer, T., Luyten, L., Aerts, J. M., & Nuttin, B. (2014). Rethinking food anticipatory activity in the activity-based anorexia rat model. Sci Rep, 4, 3929. 10.1038/srep03929

102. Xiao, H., Huang, C., Wu, Y., Wang, J. J., & Wang, H. (2025). Establishing a social behavior paradigm for female mice. Front Neurosci, 19, 1630491. 10.3389/fnins.2025.1630491

103. Yao, S., Kuja-Halkola, R., Thornton, L. M., Runfola, C. D., D’Onofrio, B. M., Almqvist, C., Lichtenstein, P., Sjölander, A., Larsson, H., & Bulik, C. M. (2016). Familial Liability for Eating Disorders and Suicide Attempts: Evidence From a Population Registry in Sweden. JAMA Psychiatry, 73(3), 284–291. 10.1001/jamapsychiatry.2015.2737

104. Zanikov, T., Gerasymchuk, M., Ghasemi Gojani, E., Robinson, G. I., Asghari, S., Groves, A., Haselhorst, L., Nandakumar, S., Stahl, C., Cameron, M., Li, D., Rodriguez-Juarez, R., Snelling, A., Hudson, D., Fiselier, A., Kovalchuk, O., & Kovalchuk, I. (2023). The Effect of Combined Treatment of Psilocybin and Eugenol on Lipopolysaccharide-Induced Brain Inflammation in Mice. Molecules, 28(6). 10.3390/molecules28062624

