## Supplementary for "Psilocybin exerts differential effects on social behaviour and inflammation in mice in contexts of activity-based anorexia (ABA)"

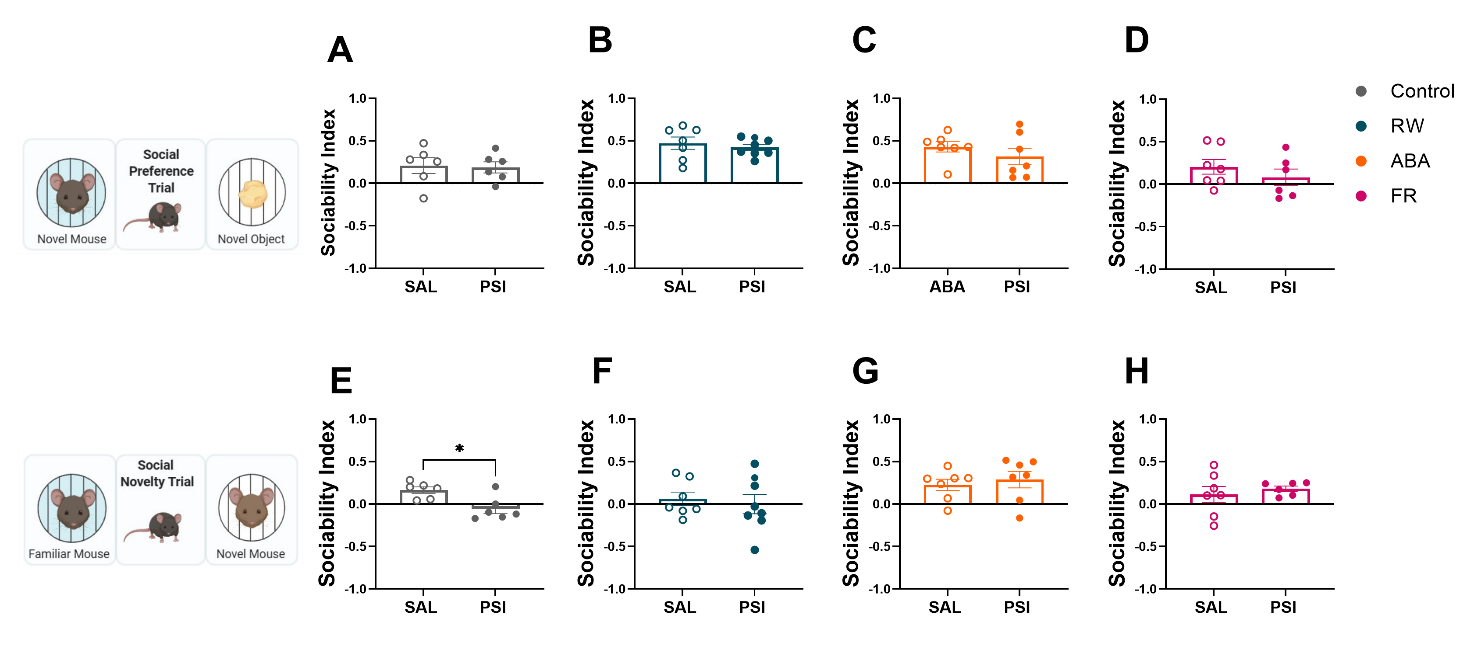

**Supplementary Data 1. Effects of a single dose of psilocybin (1.5 mg/kg) on social interaction in female mice.** During the social preference trial, PSI did not affect the preference of (A) controls, (B) RW, (C) ABA or (D) FR groups. However, during the social novelty trial, (E) PSI increased the preference of controls towards familiarity compared to novelty (p= 0.0102). PSI did not affect the social preference of (F) RW, (G) ABA or (H) FR mice during this trial. ABA (Activity-Based Anorexia), FR (food-restricted), RW (running wheel) groups; psilocybin (PSI), saline (SAL). Sociability index was based on the total time spent actively sniffing the novel mouse versus the novel object or the familiar mouse. Data are mean ± SEM, analysed by an unpaired t-test. Significance levels are indicated as: *p < 0.05

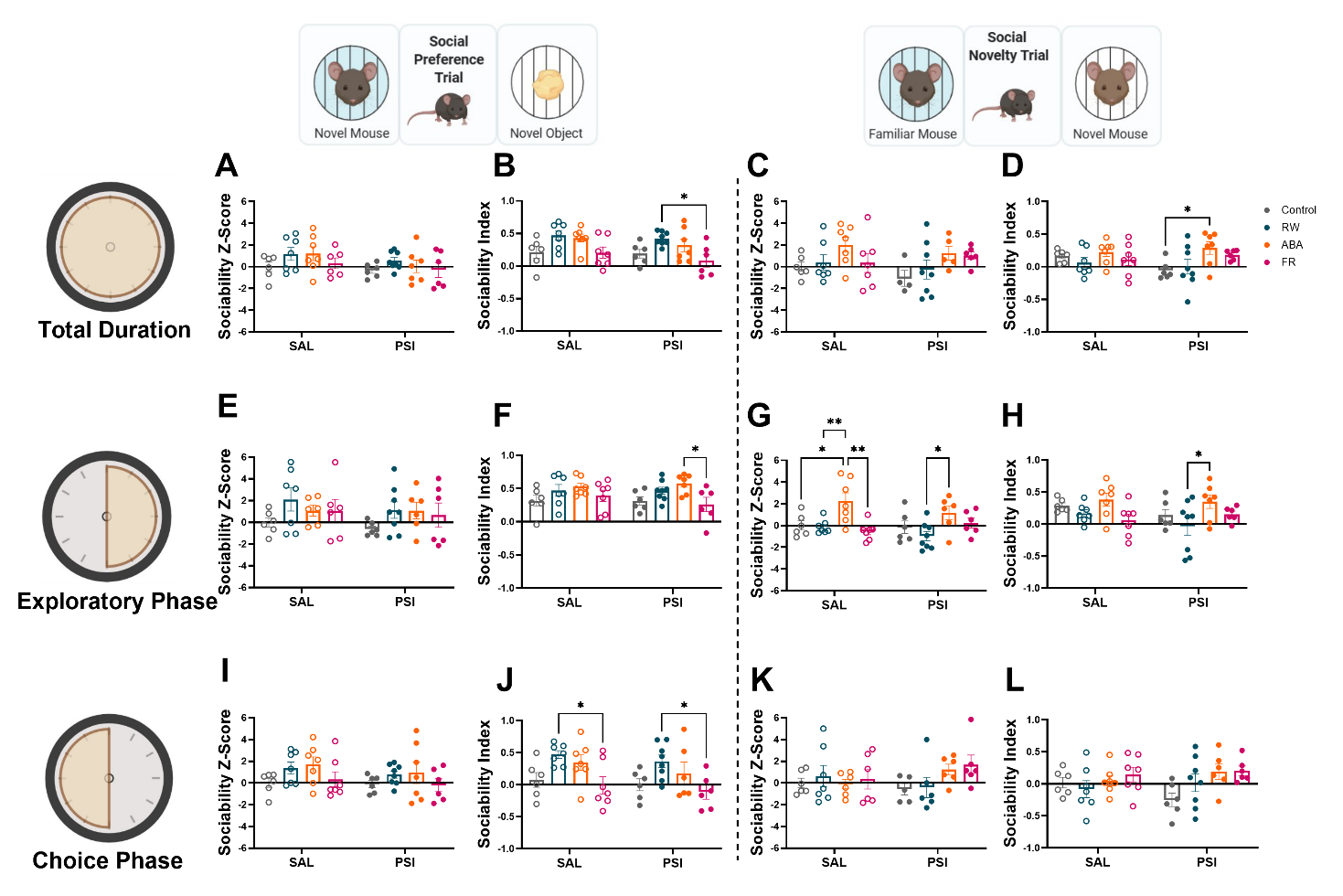

**Supplementary Data 2. Effects of a single dose of psilocybin (1.5 mg/kg) on sociability in female mice.** (A,C,E,I,K) Psilocybin treatment did not significantly affect sociability Z-scores or sociability during the total, exploratory, or choice phases of the social preference and social novelty trials. (B,D,F,G,H,J) In the Social Preference Trial, (B) PSI-treated RW mice exhibited a higher SI than FR mice (p = 0.0134). (D) PSI-treated ABA mice showed higher sociability than RW mice (p = 0.0309) during the total duration of the social novelty trial. (F) PSI-treated ABA mice had higher sociability than FR mice during the social preference trial (p = 0.0234). In the Social Novelty Trial, (G) SAL-treated ABA mice had higher sociability Z-scores than controls (p = 0.0156), RW (p = 0.007) and FR (p = 0.0012), while PSI-treated ABA mice also showed higher Z-scores than RW (p = 0.0178) during the exploratory phase of the social novelty trial. (H) Furthermore, during the same phase of the trial, PSI-treated ABA mice exhibited higher sociability than RW (p = 0.0236). (J) Lastly, during the choice phase of the social preference trial, SAL-treated RW mice had higher sociability than FR (p = 0.021), and PSI-treated RW mice had higher SI than FR (p = 0.0233). ABA (Activity-Based Anorexia), FR (food-restricted), RW (running wheel) groups; psilocybin (PSI), saline (SAL). Empty symbols represent saline-treated mice; filled symbols represent psilocybin-treated mice. Sociability Z-scores were calculated based on time spent and frequency of entries into the novel mouse versus novel object chambers, normalised to control values. Sociability was based on the total time spent actively sniffing the novel mouse versus the novel object. Data are mean ± SEM, analysed by two-way ANOVA with Tukey post hoc tests. Significance levels are indicated as: *p < 0.05; **p < 0.01; *** p < 0.001.

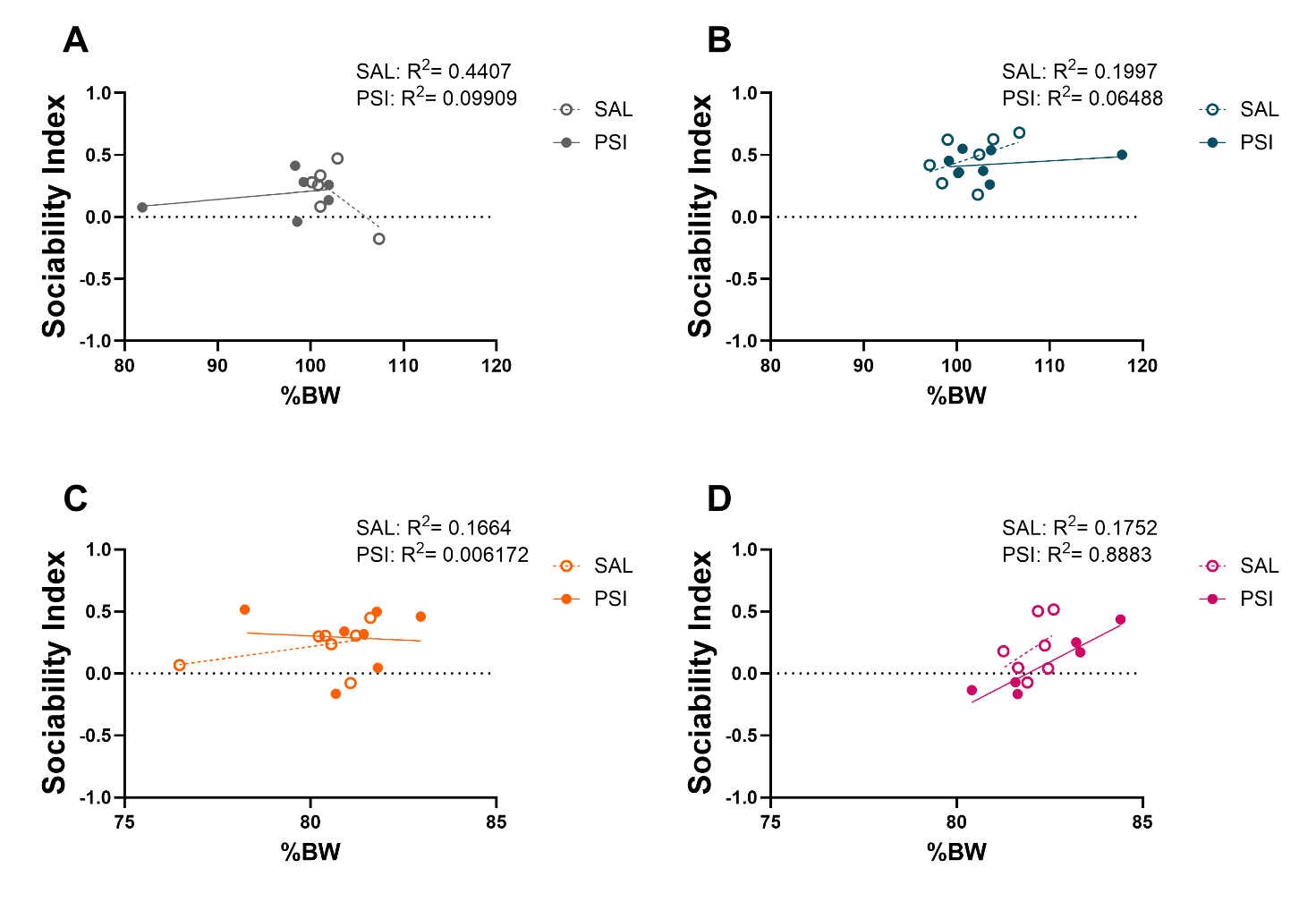

**Supplementary Data 3. Correlation between Sociability Index during the total duration of social preference trial and %BW of mice.** No correlation was observed between sociability and %BW in SAL and PSI-treated (A) Control, (B) RW and (C) ABA mice. However, PSI-treated FR mice showed a strong correlation between their sociability and %BW (Pearson's r= 0.9425, p= 0.0049) while this was not seen in the SAL-treated FR mice. ABA (Activity-Based Anorexia), FR (food-restricted), RW (running wheel) groups; psilocybin (PSI), saline (SAL). Empty symbols represent saline-treated mice; filled symbols represent psilocybin-treated mice.

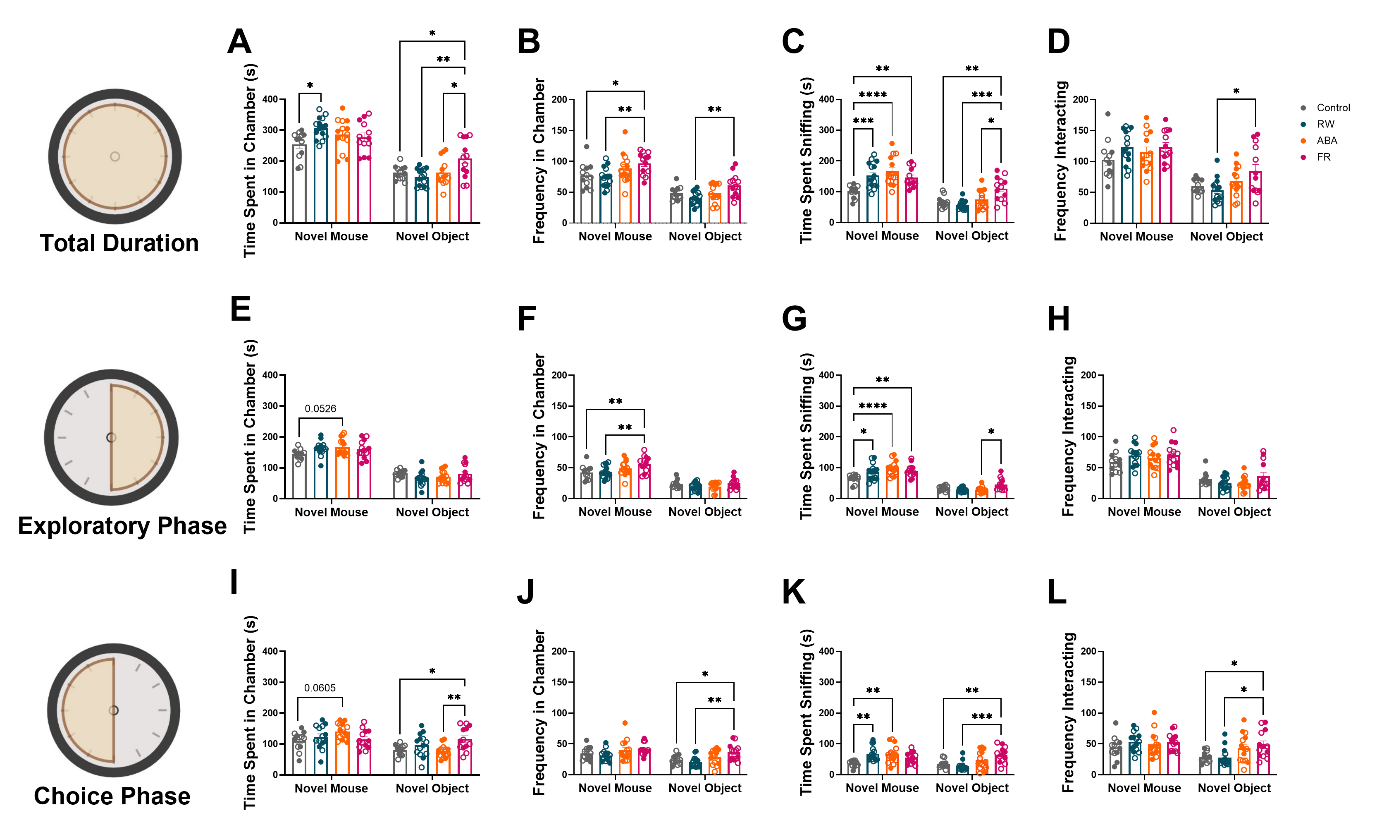

**Supplementary Data 4. Time and frequency spent exploring during social preference trial.** During the total duration of the social preference trial, (A) RW mice spent significantly more time in the novel mouse chamber compared to controls (p = 0.0103). The FR group spent significantly more time in the novel object chamber compared to controls (p = 0.0422), RW mice (p = 0.0018), and ABA mice (p = 0.0372). The FR group also showed a higher number of entries to the novel mouse chamber compared to controls (p = 0.026) and RW mice (p = 0.0042), and a higher frequency of entries into the novel object chamber compared to RW mice (p = 0.0094). (C) ABA mice spent significantly more time sniffing the novel mouse compared to controls (p < 0.0001). The FR group spent significantly more time sniffing the novel mouse compared to controls (p = 0.0044) and RW mice (p = 0.0004). A significantly higher preference towards the novel object was observed in the FR group compared to controls (p = 0.0062), RW mice (p = 0.0001), and ABA mice (p = 0.026). (D) The FR group interacted with the novel object more frequently than the RW group (p = 0.0172). During the exploratory phase of the trial, (E) a trend towards significance in the preference of ABA mice for the novel mouse compared to the control group (p = 0.0526) was observed. (F) The FR group had a higher frequency in the novel mouse chamber compared to the control group (p = 0.0052) and RW mice (p = 0.0061). (G) A significant difference in the preference towards the novel mouse was observed between control and RW groups (p = 0.0152), control group and ABA mice (p < 0.0001), and control group and FR mice (p = 0.0022). The FR group showed a significantly higher interaction time with the novel object compared to the ABA group (p = 0.0463). During the choice phase, (I) a trend towards significance in the preference of ABA mice in spending time in the chamber with the novel mouse compared to the control group (p = 0.0605). The FR group spent significantly more time in the novel object chamber compared to controls (p = 0.0191) and ABA mice (p = 0.0036). (J) The FR group had a higher frequency entering the novel object chamber compared to the control group (p = 0.0459) and RW mice (p = 0.002). A significant increase in the preference towards the novel mouse was observed in RW groups (p = 0.0022) and ABA mice (p = 0.0036) compared to controls. (K) The FR group spent significantly more time interacting with the novel object compared to controls (p = 0.0044) and RW mice (p = 0.0001). (L) The FR group had a higher frequency of sniffing the novel object compared to the control group (p = 0.0331) and RW mice (p = 0.0118). ABA (Activity-Based Anorexia), FR (food-restricted), RW (running wheel) groups. Empty symbols represent saline-treated mice; filled symbols represent psilocybin-treated mice. Data are mean ± SEM, analysed by two-way ANOVA with Tukey post hoc tests. Significance levels are indicated as: *p < 0.05; **p < 0.01; ***p < 0.001; ****p < 0.0001.

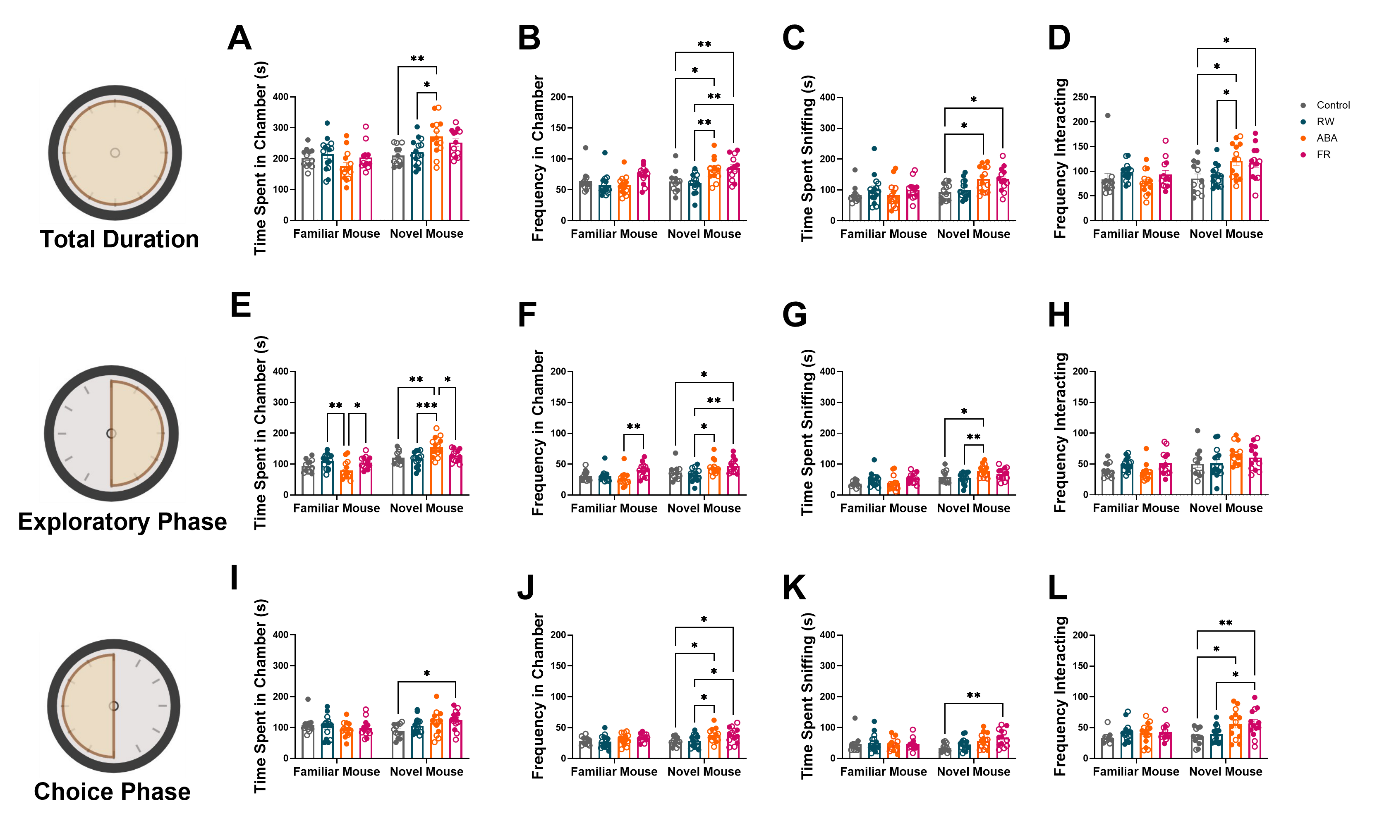

**Supplementary Data 5. Time and frequency spent exploring during social novelty trial.** During the total duration of the social novelty trial, (A) ABA mice spent significantly more time in the novel mouse chamber compared to controls (p = 0.0035) and RW mice (p = 0.0149). (B) The FR group had a higher frequency of entries to the novel mouse chamber compared to controls (p = 0.0072) and RW mice (p = 0.0011). ABA mice also had a higher frequency of entries into the novel mouse chamber compared to controls (p = 0.0206) and RW mice (p = 0.0039). (C) ABA mice spent significantly more time sniffing the novel mouse compared to controls (p = 0.0184), and FR mice spent more time sniffing the novel mouse compared to controls (p = 0.0263). (D) The ABA group interacted with the novel mouse more frequently than controls (p = 0.0147) and RW mice (p = 0.0409). The FR group interacted with the novel mouse more than controls (p = 0.0424). During the exploratory phase of the trial, (E) ABA mice exhibited a significantly lower preference for the familiar mouse compared to RW mice (p = 0.0052) and FR mice (p = 0.0329). ABA mice also showed a significantly higher preference for the novel mouse chamber compared to controls (p = 0.0022), RW mice (p = 0.0001), and FR mice (p = 0.0186). (F) The FR group had a higher frequency of exploring the novel mouse chamber compared to ABA mice (p = 0.0056), controls (p = 0.0476), and RW mice (p = 0.0016). ABA mice showed a higher number of entries in the novel mouse chamber compared to RW mice (p = 0.0117). (G) A significant difference in the time sniffing the novel mouse was observed between controls and ABA mice (p = 0.042), and ABA mice and RW mice (p = 0.0081). During the choice phase of the trial, (I) a significantly higher preference of the FR group to spend time in the novel mouse chamber compared to controls was observed (p = 0.0237). (J) The FR group had a higher frequency of entering the novel mouse chamber compared to controls (p = 0.0222) and RW mice (p = 0.0489). ABA mice entered the novel chamber more than controls (p = 0.0164) and RW mice (p = 0.0367). (K) A significant increase in the preference towards the novel mouse was observed in FR mice compared to controls (p = 0.0037). (L) The FR group had a higher frequency of sniffing the novel mouse compared to controls (p = 0.0092) and RW mice (p = 0.0405). ABA mice also sniffed the novel mouse more than controls (p = 0.013). ABA (Activity-Based Anorexia), FR (food-restricted), RW (running wheel) groups. Empty symbols represent saline-treated mice; filled symbols represent psilocybin-treated mice. Data are mean ± SEM, analysed by two-way ANOVA with Tukey post hoc tests. Significance levels are indicated as: *p < 0.05; **p < 0.01.

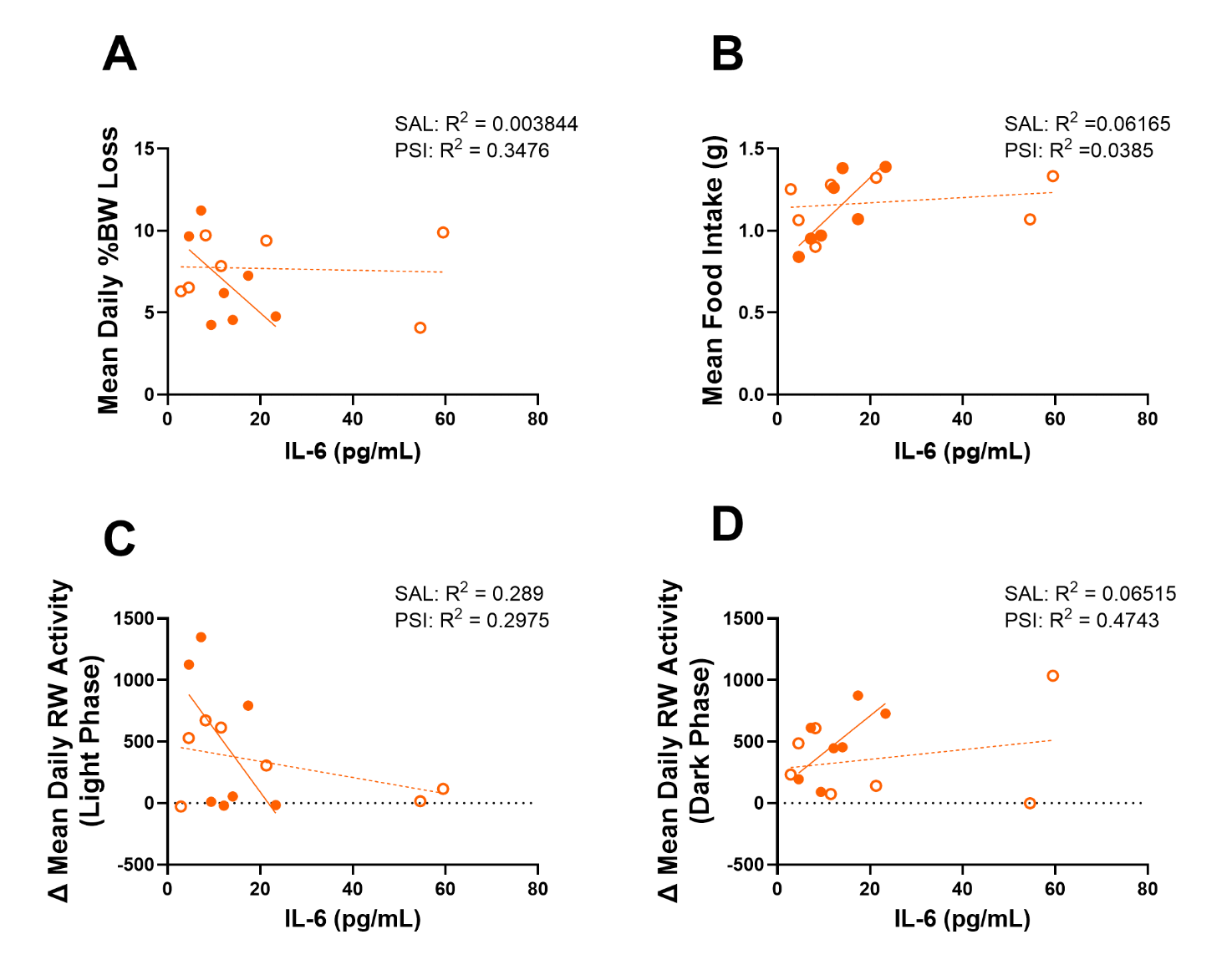

**Supplementary Data 6. Correlation between IL-6 and different parameters in ABA mice treated with psilocybin or saline control.** (A) In both SAL and psilocybin PSI-treated mice, IL-6 levels did not correlate with mean daily body weight loss. (B) a strong correlation was observed between IL-6 levels and mean food intake in PSI-treated but not SAL-treated ABA mice (Pearson r= 0.7801, p = 0.0385). No significant correlations were observed between IL-6 and (C) Δ daily light-phase or (D) dark-phase running wheel RW activity in either SAL- or PSI-treated ABA mice. Empty symbols represent saline-treated mice; filled symbols represent psilocybin-treated mice. saline (SAL), psilocybin (PSI).

### Statistics Table- Figure 1.

| **Figure** | **Statistical test** | **Group n** | **Main analysis result** | **Post-hoc multiple comparisons of interest** |
| --- | --- | --- | --- | --- |
| 1.B | Mixed-effects model | ABA (n=15), FR (n=14) | Main effect of time: F(4.536,120.2)=310.8, p<0.0001; Group × time interaction: F(4.536,120.2)=6.30, p<0.0001 | %BW: ABA < FR on Day 3 (p=0.036) and Day 4 (p=0.0263) |
| 1.C | Two-way ANOVA | ABA (n=15), FR (n=14) | Group effect: F(1,27)=8.27, p=0.0078; Period effect: F(1,27)=366.7, p<0.0001; Interaction: F(1,27)=22.99, p<0.0001 | ABA > FR, p<0.0001 |
| 1.D | Mixed-effects model | ABA (n=15), FR (n=14) | No significant effects | None |
| 1.E | Two-way ANOVA | ABA (n=15), FR (n=14) | Period: F(1,27) = 1058, p < 0.0001; Period × group interaction: F(1,27) = 8.69, p = 0.0065 | Habituation: ABA > FR (p = 0.0114) |
| 1.F | Mixed-effects model | ABA (n= 15), RW (n= 15) | Time: F(4.344,119.5) = 12.54, p < 0.0001; Group × time: F(4.344,119.5) = 3.67, p = 0.006 | Day 4: ABA < RW (p = 0.0245) |
| 1.G | Two-way ANOVA | ABA (n= 15), RW (n= 15) | Group: F(1,28) = 9.05, p = 0.0055 | Light phase: ABA > RW (p = 0.0294); Dark phase: trend (p = 0.088) |
| 1.H | Two-way ANOVA | ABA (n= 15), RW (n= 15) | Time: F(23,644) = 17.94, p < 0.0001; Group: F(1,28) = 8.60, p = 0.0066; Group × time: F(23,644) = 9.43, p < 0.0001 | ABA < RW: 0900h (p = 0.0477), 1100h (p = 0.0176), 0800h (p = 0.01); ABA > RW: 1600h (p = 0.0006), 1700h (p < 0.0001), 1800h (p = 0.0005), 0300h (p = 0.0043), 0400h (p = 0.0018), 0600h (p = 0.0026), 0700h (p = 0.0001) |
| 1.I | Two-way ANOVA | ABA (n= 15), RW (n= 15) | Time: F(23,644) = 13.98, p < 0.0001; Group × time: F(23,644) = 7.68, p < 0.0001 | ABA < RW: 1000h (p = 0.001), 1100–1200h (p < 0.0001), 0800h (p = 0.0004); ABA > RW: 1600h (p = 0.0136), 1700h (p = 0.0301) |

### Statistics Table- Figure 2.

| **Figure** | **Statistical test** | **Group n** | **Main analysis result** | **Post-hoc multiple comparisons of interest** |
| --- | --- | --- | --- | --- |
| 2A | Unpaired t-test | ABA (n= 15), RW (n= 15) | No significant group differences | None |
| 2B | Two-way ANOVA | ABA (n= 15), RW (n= 15) | Period effect: F(1,28) = 20.71, p < 0.0001 | None |
| 2C | Two-way ANOVA | ABA (n= 15), RW (n= 15) | Period effect: F(1,28) = 19.51, p = 0.0001; Period × group interaction: F(1,28) = 5.663, p = 0.0244 | ABA > RW during ABA period (p = 0.0191) |
| 2D | Two-way ANOVA | ABA (n= 15), RW (n= 15) | Period effect: F(1,28) = 40.56, p < 0.0001; Period × group interaction: F(1,28) = 4.761, p = 0.0377 | None |
| 2E | Mixed-effects model | ABA (n= 15), RW (n= 15) | Time effect: F(263,7244) = 44.66, p < 0.0001 | None |
| 2F | Pearson correlation | ABA (n= 15), RW (n= 15) | ABA: r = 0.7251, p = 0.0022 |  |
| 2G | Pearson correlation | ABA (n= 15), RW (n= 15) | No significant correlation |  |

### Statistics Table- Figure 3.

| **Figure** | **Statistical test** | **Group n** | **Main analysis result** | **Post-hoc multiple comparisons of interest** |
| --- | --- | --- | --- | --- |
| 3.A | One-way ANOVA | Control (n= 12), FR (n= 13), ABA (n= 14), RW (n= 15) | No significant group differences | None |
| 3.B | One-way ANOVA | Control (n= 12), FR (n= 13), ABA (n= 14), RW (n= 15) | No significant group differences | None |
| 3.C | One-way ANOVA | Control (n= 12), FR (n= 13), ABA (n= 14), RW (n= 15) | Group effect: F(3,50) = 7.05, p = 0.0005 | RW > Control (p = 0.0123); RW > FR (p = 0.0012); ABA > FR (p = 0.0278) |
| 3.D | One-way ANOVA | Control (n= 12), FR (n= 12), ABA (n= 12), RW (n= 15) | No significant group differences | None |
| 3.E | One-way ANOVA | Control (n= 12), FR (n= 13), ABA (n= 14), RW (n= 15) | Group effect: F(3,50) = 4.93, p = 0.0045 | ABA > Control (p = 0.0136); ABA > FR (p = 0.020) |
| 3.F | One-way ANOVA | Control (n= 12), FR (n= 13), ABA (n= 13), RW (n= 15) | No significant group differences | None |
| 3.G | One-way ANOVA | Control (n= 12), FR (n= 13), ABA (n= 14), RW (n= 15) | Group effect: F(3,49) = 7.44, p = 0.0003 | RW > Control (p = 0.0102); RW > FR (p = 0.0005); ABA > FR (p = 0.0366) |

### Statistics Table- Figure 4.

| **Figure** | **Statistical test** | **Group n** | **Main analysis result** | **Post-hoc multiple comparisons of interest** |
| --- | --- | --- | --- | --- |
| 4.A | One-way ANOVA | Control (n= 12), FR (n= 13), ABA (n= 14), RW (n= 15) | No significant group differences | None |
| 4.B | One-way ANOVA | Control (n= 10), FR (n= 13), ABA (n= 12), RW (n= 15) | Group effect: F(3,46) = 3.03, p = 0.039 | ABA > Control (p = 0.0209) |
| 4.C | One-way ANOVA | Control (n= 12), FR (n= 13), ABA (n= 14), RW (n= 15) | Group effect: F(3,50) = 3.23, p = 0.030 | ABA > RW (p = 0.0181); ABA vs Control (trend, p = 0.0606) |
| 4.D | One-way ANOVA | Control (n= 12), FR (n= 13), ABA (n= 13), RW (n= 15) | Group effect: F(3,49) = 8.42, p = 0.0001 | ABA > Control (p = 0.0059); ABA > RW (p = 0.0027); ABA > FR (p = 0.0001) |
| 4.E | One-way ANOVA | Control (n= 12), FR (n= 13), ABA (n= 14), RW (n= 15) | Group effect: F(3,50) = 4.17, p = 0.010 | ABA > RW (p = 0.0112); ABA vs FR (trend, p = 0.0537) |
| 4.F | One-way ANOVA | Control (n= 12), FR (n= 12), ABA (n= 13), RW (n= 14) | No significant group differences | None |
| 4.G | One-way ANOVA | Control (n= 12), FR (n= 13), ABA (n= 13), RW (n= 15) | No significant group differences | None |

### Statistics Table- Figure 5.

| **Figure** | **Statistical test** | **Group n** | **Main analysis result** | **Post-hoc multiple comparisons of interest** |
| --- | --- | --- | --- | --- |
| 5.A | One-way ANOVA | Control (n= 12), FR (n= 14), ABA (n= 14), RW (n= 12) | No significant group differences | None |
| 5.B | Two-way ANOVA | Control-SAL (n= 6), Control-PSI (n= 6), FR-SAL (n= 7), FR-PSI (n= 7), ABA-SAL (n= 7), ABA-PSI (n= 7), RW-SAL (n= 7), RW-PSI (n= 6) | Main effect of group: F(3,44) = 3.024, p = 0.0395; Treatment × group interaction: F(3,44) = 3.637, p = 0.0198 | RW- PSI > RW- SAL (p = 0.0029); RW- PSI > Controls- PSI (p = 0.005); RW- PSI > ABA- PSI (p = 0.011) |
| 5.C | Pearson correlation | Control-SAL (n= 6), Control-PSI (n= 6), RW-SAL (n= 7), RW-PSI (n= 6) | RW- PSI: r = 0.8020, p = 0.0166; Control- PSI: r = 0.8751, p = 0.0224 |  |

### Statistics Table- Supplementary Data 1.

| **Figure** | **Statistical test** | **Group n** | **Main analysis result** | **Post-hoc multiple comparisons of interest** |
| --- | --- | --- | --- | --- |
| S1A | t-test | Control- SAL (n= 6), Control- PSI (n= 6) | No significant group differences | None |
| S1B | t-test | RW- SAL (n= 7), RW- PSI (n= 8) | No significant group differences | None |
| S1C | t-test | ABA- SAL (n= 7), ABA- PSI (n= 7) | No significant group differences | None |
| S1D | t-test | FR- SAL (n= 7), FR- PSI (n= 6) | No significant group differences | None |
| S1E | t-test | Control- SAL (n= 6), Control- PSI (n= 6) | t(10)= 3.16, p= 0.0102 |  |
| S1F | t-test | RW- SAL (n= 7), RW- PSI (n= 8) | No significant group differences | None |
| S1G | t-test | ABA- SAL (n= 7), ABA- PSI (n= 7) | No significant group differences | None |
| S1H | t-test | FR- SAL (n= 7), FR- PSI (n= 6) | No significant group differences | None |

### Statistics Table- Supplementary Data 2.

| **Figure** | **Statistical test** | **Group n** | **Main analysis result** | **Post-hoc multiple comparisons of interest** |
| --- | --- | --- | --- | --- |
| S2A, C, E, I, K, L | Two-way ANOVA | Control-SAL (n= 6), Control-PSI (n= 6), FR-SAL (n= 7), FR-PSI (n= 7), ABA-SAL (n= 7), ABA-PSI (n= 7), RW-SAL (n= 7), RW-PSI (n= 6) | No significant treatment effects | None |
| S2B | Two-way ANOVA | Control-SAL (n= 6), Control-PSI (n= 6), FR-SAL (n= 7), FR-PSI (n= 7), ABA-SAL (n= 7), ABA-PSI (n= 7), RW-SAL (n= 7), RW-PSI (n= 6) | Group effect: F(3,46) = 7.05, p = 0.0005 | RW-PSI > FR-PSI (p = 0.0134) |
| S2D | Two-way ANOVA | Control-SAL (n= 6), Control-PSI (n= 6), RW-SAL (n= 7), RW-PSI (n= 6) | Group effect: F(3,46) = 3.20, p = 0.032 | ABA-PSI > Control-PSI (p = 0.0309) |
| S2F | Two-way ANOVA | Control, FR, ABA, RW; Sal vs PSI | Group effect: F(3,46) = 4.81, p = 0.0054 | ABA-PSI > FR-PSI (p = 0.0234) |
| S2G | Two-way ANOVA | Control-SAL (n= 6), Control-PSI (n= 6), FR-SAL (n= 7), FR-PSI (n= 7), ABA-SAL (n= 7), ABA-PSI (n= 7), RW-SAL (n= 7), RW-PSI (n= 6) | Group effect: F(3,45) = 8.05, p = 0.0002 | ABA- SAL > Control- SAL (p = 0.0156), RW- SAL (p = 0.007), FR- SAL (p = 0.0012); ABA- PSI > RW-PSI (p = 0.0178) |
| S2H | Two-way ANOVA | Control-SAL (n= 6), Control-PSI (n= 6), FR-SAL (n= 7), FR-PSI (n= 7), ABA-SAL (n= 7), ABA-PSI (n= 7), RW-SAL (n= 7), RW-PSI (n= 6) | Group effect: F(3,46) = 4.00, p = 0.013 | ABA- PSI > RW-PSI (p = 0.0236) |
| S2J | Two-way ANOVA | Control-SAL (n= 6), Control-PSI (n= 6), FR-SAL (n= 7), FR-PSI (n= 7), ABA-SAL (n= 7), ABA-PSI (n= 7), RW-SAL (n= 7), RW-PSI (n= 6) | Group effect: F(3,45) = 7.25, p = 0.0005 | RW-PSI > FR-PSI (p = 0.021); RW-PSI > FR-PSI (p = 0.0233) |

### Statistics Table- Supplementary Data 3.

| **Figure** | **Statistical test** | **Group n** | **Main analysis result** |
| --- | --- | --- | --- |
| S3A | Pearson correlation | Control-SAL (n= 6), Control-PSI (n=6) | No significant correlation |
| S3B | Pearson correlation | RW-SAL (n= 7), RW-PSI (n= 8) | No significant correlation |
| S3C | Pearson correlation | ABA-SAL (n= 7), ABA-PSI (n= 7) | No significant correlation |
| S3D | Pearson correlation | FR-SAL (n= 7), FR-SAL (n= 6) | FR-PSI: Pearson's r= 0.9425, p= 0.0049 |

### Statistics Table- Supplementary Data 4.

| **Figure** | **Statistical test** | **Group n** | **Main analysis result** | **Post-hoc multiple comparisons of interest** |
| --- | --- | --- | --- | --- |
| S4A-D | Two-way ANOVA | Control (n= 12), FR (n= 13), ABA (n= 14), RW (n= 15) | Chamber effects: time F(1,100)=177, p<0.0001; entries F(1,100)=112, p<0.0001; sniffing time F(1,100)=107, p<0.0001; interaction freq F(1,100)=88.94, p<0.0001. Group effects: entries F(3,100)=8.008, p<0.0001; sniffing time F(3,100)=9.108, p<0.0001; object interaction F(3,100)=2.952, p=0.0363. Interactions: chamber × group (time F(3,100)=5.851, p=0.001; sniffing F(3,100)=7.667, p<0.0001). | RW > Control (time in novel mouse chamber, p=0.0103). FR > Control, RW, ABA (time in novel object chamber, p=0.0422, p=0.0018, p=0.0372). FR > Control, RW (entries into novel mouse, p=0.026, p=0.0042). FR > RW (entries into novel object, p=0.0094). ABA > Control (time sniffing novel mouse, p<0.0001). FR > Control, RW (time sniffing novel mouse, p=0.0044, p=0.0004). FR > Control, RW, ABA (novel object preference, p=0.0062, p=0.0001, p=0.026). FR > RW (object interaction freq, p=0.0172). |
| S4E-H | Two-way ANOVA | Control (n= 12), FR (n= 13), ABA (n= 14), RW (n= 15) | Chamber effects: time F(1,100)=343.7, p<0.0001; sniff freq F(1,100)=194.6, p<0.0001; sniff time F(1,100)=225.1, p<0.0001; object sniff freq F(1,100)=146.5, p<0.0001. Group effects: sniff freq F(3,100)=4.131, p=0.0083; sniff time F(3,100)=5.701, p=0.0012. Interactions: chamber × group (time F(3,100)=3.29, p=0.0238; sniff time F(3,100)=6.056, p=0.0008). | ABA vs Control (novel mouse preference trend, p=0.0526). FR > Control, RW (freq novel mouse chamber, p=0.0052, p=0.0061). FR > Control, ABA (time in novel object chamber, p=0.0191, p=0.0036). Controls < RW, ABA, FR (novel mouse preference, p=0.0152, p<0.0001, p=0.0022). FR > ABA (novel object interaction time, p=0.0463). |
| S4I-L | Two-way ANOVA | Control (n= 12), FR (n= 13), ABA (n= 14), RW (n= 15) | Chamber effects: time F(1,100)=25.9, p<0.0001; sniff freq F(1,100)=17.81, p<0.0001; sniff time F(1,100)=10.08, p=0.002; object sniff freq F(1,100)=13.48, p=0.0004. Group effects: sniff freq F(3,100)=6.807, p=0.0003; sniff time F(3,100)=6.639, p=0.0005; object sniff freq F(3,100)=3.174, p=0.0275. Interactions: chamber × group (time F(3,100)=5.621, p=0.0013; sniff time F(3,100)=6.683, p=0.0004). | ABA vs Control (novel mouse preference trend, p=0.0605). FR > Control, ABA (time in novel object chamber, p=0.0191, p=0.0036). FR > Control, RW (entries into novel object, p=0.0459, p=0.002). RW, ABA > Control (novel mouse preference, p=0.0022, p=0.0036). FR > Control, RW (object interaction time, p=0.0044, p=0.0001). FR > Control, RW (object sniff freq, p=0.0331, p=0.0118). |

### Statistics Table- Supplementary Data 5.

| **Figure** | **Statistical test** | **Group n** | **Main analysis result** | **Post-hoc multiple comparisons of interest** |
| --- | --- | --- | --- | --- |
| S5A-D | Two-way ANOVA | Control (n= 12), FR (n= 13), ABA (n= 14), RW (n= 15) | Chamber effects: time F(1,100)=21.79, p<0.0001; entries F(1,100)=9.108, p=0.0032; sniffing time F(1,100)=11.86, p=0.0008; interactions F(1,100)=7.467, p=0.0074. Group effects: entries F(3,100)=7.615, p=0.0001; sniffing time F(3,100)=3.034, p=0.0327; interactions F(3,100)=4.184, p=0.0078. Interactions: chamber × group (time F(3,100)=6.317, p=0.0006; entries F(1,100)=3.149, p=0.0283; sniffing F(3,100)=2.942, p=0.0367). | ABA > Control, RW (time in novel mouse chamber, p=0.0035, p=0.0149). FR > Control, RW (entries into novel mouse chamber, p=0.0072, p=0.0011). ABA > Control, RW (entries into novel mouse chamber, p=0.0206, p=0.0039). ABA > Control (sniffing novel mouse, p=0.0184). FR > Control (sniffing novel mouse, p=0.0263). ABA > Control, RW (frequency novel mouse interaction, p=0.0147, p=0.0409). FR > Control (frequency novel mouse interaction, p=0.0424). |
| S5E-H | Two-way ANOVA | Control (n= 12), FR (n= 13), ABA (n= 14), RW (n= 15) | Chamber effects: time F(1,100)=49.06, p<0.0001; sniff freq F(1,100)=14.72, p=0.0002; entries F(1,99)=28.79, p<0.0001; sniff novel mouse/object F(1,100)=13.72, p=0.0003. Group effects: sniff freq F(3,100)=6.974, p=0.0003; entries F(3,99)=4.207, p=0.0076. Interactions: chamber × group (time F(3,100)=11.23, p<0.0001; sniff freq F(1,100)=3.214, p=0.0261; sniff novel mouse/object F(1,100)=3.001, p=0.0341). | ABA < RW, FR (preference for familiar mouse, p=0.0052, p=0.0329). ABA > Control, RW, FR (preference for novel mouse chamber, p=0.0022, p=0.0001, p=0.0186). FR > ABA, Control, RW (freq exploring novel mouse chamber, p=0.0056, p=0.0476, p=0.0016). ABA > RW (entries novel mouse chamber, p=0.0117). ABA > Control (novel mouse preference, p=0.042). ABA > RW (novel mouse preference, p=0.0081). |
| S5I-L | Two-way ANOVA | Control (n= 12), FR (n= 13), ABA (n= 14), RW (n= 15) | Chamber effects: time F(1,100)=25.9, p<0.0001; sniff freq F(1,100)=17.81, p<0.0001; sniff time F(1,100)=10.08, p=0.002; sniff novel mouse/object F(1,100)=13.48, p=0.0004. Group effects: entries F(3,100)=5.648, p=0.0013; sniff time F(3,100)=6.639, p=0.0005; sniff novel mouse/object F(3,100)=3.174, p=0.0275. Interactions: chamber × group (time F(3,100)=2.95, p=0.0364; sniff time F(3,100)=2.818, p=0.0429). | FR > Control (time in novel mouse chamber, p=0.0237). FR > Control, RW (entries novel mouse chamber, p=0.0222, p=0.0489). ABA > Control, RW (entries novel mouse chamber, p=0.0164, p=0.0367). ABA > Control (novel mouse preference, p=0.0037). FR > Control, RW (sniffing novel mouse, p=0.0092, p=0.0405). ABA > Control (sniffing novel mouse, p=0.013). |

### Statistics Table- Supplementary Data 6.

| **Figure** | **Statistical test** | **Group n** | **Main analysis result** |
| --- | --- | --- | --- |
| S6A | Pearson correlation | ABA-SAL (n= 7), ABA-PSI (n= 7) | No significant correlation |
| S6B | Pearson correlation | ABA-SAL (n= 7), ABA-PSI (n= 7) | SAL: No significant correlation  PSI: r = 0.7801, p = 0.0385 |
| S6C | Pearson correlation | ABA-SAL (n= 7), ABA-PSI (n= 7) | No significant correlation |
| S6D | Pearson correlation | ABA-SAL (n= 7), ABA-PSI (n= 7) | No significant correlation |
